# “Neighbourhood watch” model: embryonic epiblast cells assess positional information in relation to their neighbours

**DOI:** 10.1101/2021.04.22.440894

**Authors:** Hyung Chul Lee, Cato Hastings, Nidia M.M. Oliveira, Rubén Pérez-Carrasco, Karen M. Page, Lewis Wolpert, Claudio D. Stern

## Abstract

In many developing and regenerating systems, tissue pattern is established through gradients of informative morphogens, but we know little about how cells interpret these. Using experimental manipulation of early chick embryos including misexpression of an inducer (VG1 or ACTIVIN) and an inhibitor (BMP4), we test two alternative models for their ability to explain how the site of primitive streak formation is positioned relative to the rest of the embryo. In one model, cells read morphogen concentrations cell-autonomously. In the other, cells sense changes in morphogen status relative to their neighbourhood. We find that only the latter model can account for the experimental results, including some counter-intuitive predictions. This mechanism (which we name “neighbourhood watch” model) illuminates the classic “French Flag Problem” and how positional information is interpreted by a sheet of cells in a large developing system.

**Summary statement:** In a large developing system, the chick embryo before gastrulation, cells interpret gradients of positional signals relative to their neighbours to position the primitive streak, establishing bilateral symmetry.

## Introduction

In the late 1960s, Lewis Wolpert introduced the concept of “positional information”, asking the question of how cells within a morphogenetic field could adopt several cell-type identities in response to signalling cues from the embryo. The analogy of a French flag, with three colours: red, white and blue, was used to symbolise the cell types (Wolpert, 1968, Wolpert, 1969). Wolpert proposed that a gradient of a hypothetical “morphogen” diffusing away from a local source and decaying with distance would be read by cells, which respond with discrete thresholds to adopt the various identities. He named this the “French Flag problem”.

Since, several systems have been found in which a morphogen imparts positional information resulting in a defined morphological pattern. These include head and foot formation in Hydra (Schaller, 1973, Bode, 2011), patterning of the wing (Lecuit et al., 1996), leg and antenna imaginal discs of the fly (Postlethwait and Schneiderman, 1971) and limb regeneration in vertebrates (Kumar et al., 2007). Various mechanisms have been studied by which cells might interpret such morphogen gradients so that their positions are defined precisely and robustly. In cultured cells and explant systems (Gurdon et al., 1999, Gurdon et al., 1995) it has been shown that cells respond directly to morphogen concentration, in a manner most similar to that described by Wolpert (Wolpert, 1969). In vertebrate neural tube patterning, the gradient of Shh is transformed into a dynamic profile of Gli (a transcription factor) to generate spatial patterns of downstream gene expression, suggesting that cells interpret positional information using intracellular regulatory networks, where a temporal element is important (Dessaud et al., 2010, Cohen et al., 2013). The *bicoid* gradient, which sets up the anterior–posterior axis in fruit fly embryos has been studied extensively (Driever and Nusslein-Volhard, 1988b, Driever and Nusslein-Volhard, 1988a, Gregor et al., 2007b, Gregor et al., 2007a) and it has been suggested that spatial averaging across nuclei is one mechanism by which noise is reduced in the transduction of the *bicoid* signal (Gregor et al., 2007a).

All the above systems are relatively small (<100 cell diameters) (Wolpert, 1969) allowing stable gradients to be set up which span the entire field. However, some developing systems are much larger in size, begging the question of what mechanisms cells might use to assess their positions. An example of such a large system is the early chick embryo just before the onset of gastrulation. The embryo contains as many as 20,000-50,000 cells and is approximately 3mm in diameter. Within this large field the primitive streak, the site of gastrulation, can arise at any point around the circumference. Any isolated fragment of this large embryo can initiate primitive streak formation; however, only one primitive streak forms, suggesting the existence of patterning events that coordinate cell behaviours across the whole field.

In these early embryos, the “pattern” is established in the marginal zone, a ring-like region of extraembryonic tissue, lying just outside of the central disk-like area pellucida, where the embryo will arise. The primitive streak, the first indication of the future midline of the embryo, arises at one edge of the inner area pellucida, adjacent to the posterior part of the marginal zone, where the TGFβ-related signalling molecule cVG1 is expressed. Previous studies have shown that positioning of the primitive streak requires “positive” inducing signals by cVG1/NODAL from the posterior marginal zone near the site of streak formation, and that this is antagonised by BMP signalling which is highest at the opposite (anterior) end of the blastoderm (Fig. S1A) (Shah et al., 1997, Streit et al., 1998, Bertocchini and Stern, 2012, Streit and Stern, 1999, Bertocchini and Stern, 2002, Skromne and Stern, 2002, Bertocchini et al., 2004). The distance between the two extremes of the marginal zone is approximately 300 cell diameters. Previous studies suggested that these signals are part of a “global positioning system” to establish polarity in the chick embryo, (Bertocchini and Stern, 2012, Arias et al., 2017), and therefore that the whole embryo is a coordinated system of positional information.

To find out how cells interpret morphogen concentrations to generate positional information, we designed two computational models to represent respectively a fixed gradient, read locally by cells, or a system where cells compare themselves to their neighbours to determine their position in the field. Using a combination of embryological manipulations and computational modelling, we ask which of these two models can best account for the results of various manipulations in the spatial distribution, number and intensity of the inducing (cVG1/NODAL) and inhibitory (BMP) signals. We find that the “positional information” that determines the site of primitive streak formation is explained better by a mechanism by which cells compare themselves to their neighbours rather than by a cell autonomous assessment of gradients. We name this the “neighbourhood watch” model.

## Results

### Epiblast cells may sense local differences in strength of inducing signal rather than the absolute amount of inducer

When a small pellet of cVG1-expressing cells (HEK293T cells transfected with a cVG1-expression construct) is grafted into the anterior marginal zone (the innermost extraembryonic epiblast, just outside the central embryonic area pellucida), it can initiate formation of an ectopic primitive streak that eventually develops into a full embryonic axis (Shah et al., 1997, Skromne and Stern, 2002). However endogenous *cVG1* mRNA is expressed as a crescent encompassing an arc of about 60° in the posterior marginal zone (Fig. S1A). To mimic this distribution more closely, as well as to test the effects of greater concentration of cVG1 inducing signal, we placed two cVG1-expressing cell pellets side-by-side in the anterior marginal zone, and assessed primitive streak formation by in situ hybridisation for expression of *BRACHYURY* (*cBRA, =TBXT)* after overnight culture (Fig. 1 A-D). Only a single ectopic primitive streak was generated near the middle of the two cVG1 pellets (Fig. 1 B); neither double nor thicker ectopic streaks were observed, similar to the effects of implanting a single pellet.

**Fig. 1.**
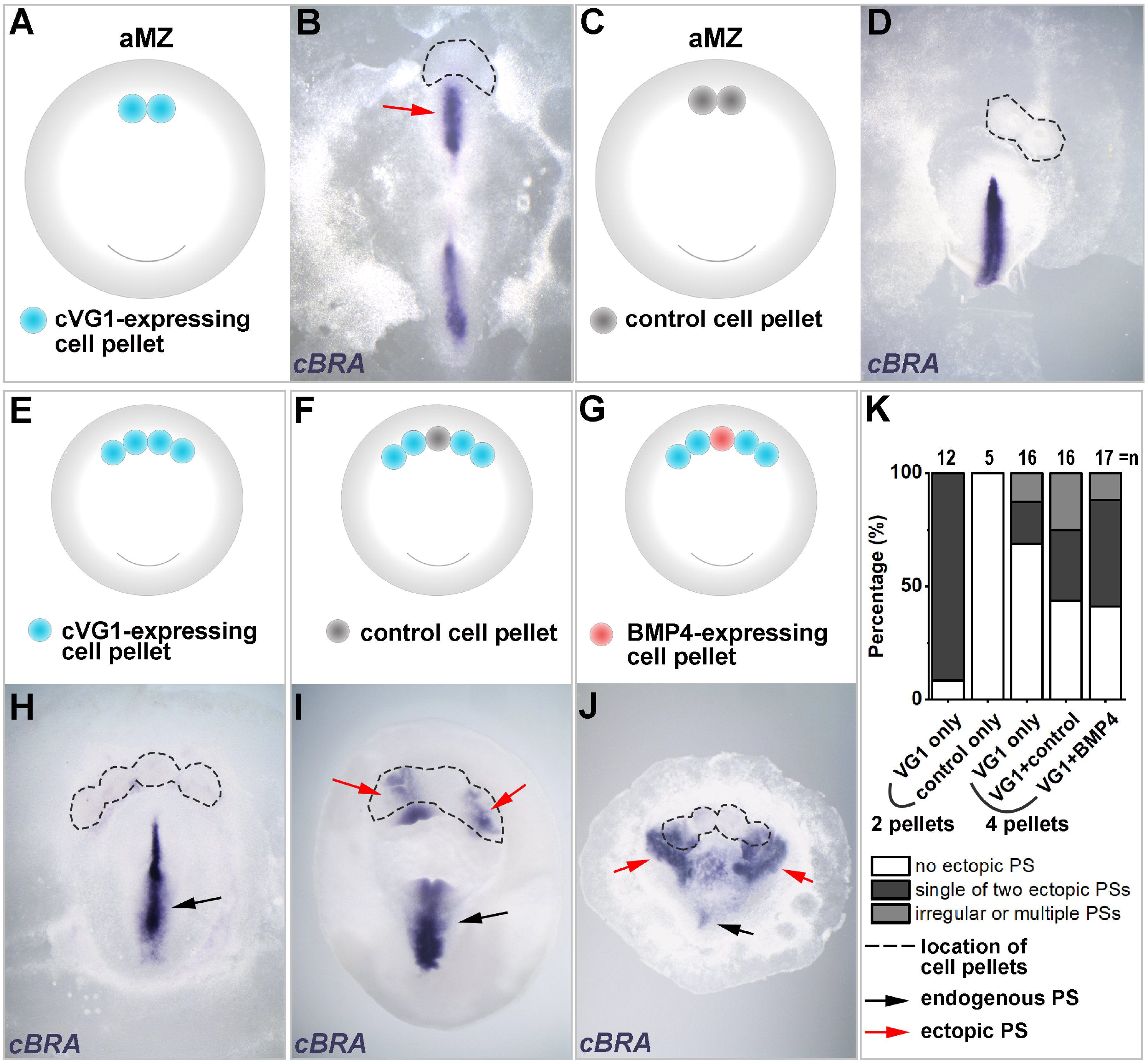
Interruption of a domain exposed to an inducing signal increases the incidence of primitive streak induction – experiments with secreting cells. **(A, B)** When two pellets of cVG1-expressing cells are grafted in the anterior marginal zone (aMZ), only a single ectopic primitive streak (red arrow) is generated. **(C, D)** Control cell pellets do not induce a streak. **(E-G)** Experimental design. Ectopic streak formation is checked in three different conditions: misexpression of cVG1 in a wide area using four cVG1-expressing cell pellets **(E)**, introduction of a ‘spacer’ (control cell pellet) to interrupt a set of four cVg1 pellets **(F)**, and introduction of an inhibitor (BMP4-expressing cell pellet) to interrupt a set of four cVg1 inducing pellets **(G)**. **(H-J)** representative embryos for each experiment. The frequency of primitive streak formation is enhanced by interrupting the domain of inducing signal, even when this interruption is achieved by introduction of an inhibitor (**J**). **(K)** Summary graph showing the incidence of each type of result for the above experiments (**E-J**). PS: primitive streak. Black and red arrows, endogenous and ectopic streaks, respectively. Dotted lines, position of the cell pellets. *cBRA*, primitive streak marker.

To provide a stronger and wider signal, we tested the effect of implanting four cVG1-expressing cell pellets side-by-side in the anterior marginal zone. Surprisingly, in the majority of cases (11/16 embryos), no ectopic primitive streak formed and no ectopic *cBRA* expression was seen (Fig. 1 E, H, K). Since application of the equivalent of a quad-dose of inducer spread over a four-fold wider area does not cause either more efficient or wider induction than a single dose, we speculated that “boundaries” to the signalling domain may be required. To test this, we placed a control cell pellet (HEK293T cells transfected with pCAβ-GFP; see Methods) to split four cVG1-expressing cell pellets into two groups on either side. The incidence of ectopic streak formation doubled (Fig. 1 F, I, K). If a boundary is indeed important, we might expect that, perhaps paradoxically, ectopic streak induction might increase if a pellet expressing the inhibitor BMP4 (rather than a control pellet) is used to interrupt the set of four cVG1-expressing cell pellets. This is indeed the case (Fig. 1 G, J, K). Together, these results suggest that cells may sense variations in signal strength in relation to their neighbours, rather than measuring the absolute amount of local signal they receive, to determine the outcome of the inductive event.

The above experiments were done using pellets of transfected cells, as in previous studies (Bertocchini et al., 2004, Bertocchini and Stern, 2012, Shah et al., 1997, Skromne and Stern, 2002, Streit et al., 1998, Torlopp et al., 2014). One problem with this approach is that cells are likely to express other (unknown) factors that could influence the outcome of the signalling event. Another problem is that these pellets are relatively large (500-1000 cells). We therefore decided to substitute the use of cell pellets with protein-soaked microbeads (about 100 μm diameter). As neither VG1 nor NODAL are available as pure proteins, we decided to use ACTIVIN instead, which can induce axial structures and mesendodermal markers in chick epiblast (Mitrani et al., 1990, Stern et al., 1995). As shown in amphibian animal cap ectoderm explants (Green and Smith, 1990), ACTIVIN also acts through the SMAD 2/3 pathway and generates finely graded responses of mesendoderm induction to different concentrations (Stern et al., 1995). BMP4-soaked beads were used as a source of inhibitory signal. First, we checked if a single soaked bead can mimic the effects of a single cell pellet (Fig. S2). Grafting a bead soaked in ACTIVIN into the anterior marginal zone has the same effect as a cell pellet placed in the same position: it induces an ectopic *cBRA*-expressing primitive streak in adjacent epiblast (Fig. S2, A-E). Conversely, placing a bead of the inhibitor BMP4 in the posterior marginal zone results in either displacement of the endogenous primitive streak to a more lateral position, or two primitive streaks, arising either side of the BMP4-bead (Fig. S2, F-J). With a high concentration of BMP4 (50 ng/μl) primitive streak formation was abolished in about half of the embryos (Fig. S2J).

Next, we mirrored the experiments done with two or more cell pellets but using soaked beads (Fig. 2). After grafting a single ACTIVIN protein-soaked bead flanked by two control beads, 43% of embryos (6/14) had ectopic *cBRA* expression (Fig. 2, B, G, K). When three ACTIVIN beads were grafted in a row to expose a wide domain to the inducing signal, the majority of embryos (78%, 7/9) showed no ectopic *cBRA* expression (Fig. 2, C, H, K). When boundaries to the signalling domain were generated either by introducing a BSA-soaked control bead (Fig.2, D and I) or a BMP4-soaked bead (Fig. 2, E and J) among the ACTIVIN beads, the proportion of embryos with ectopic *cBRA* expression was restored, to 40% (4/10) and 50% (6/12) respectively (Fig. 2 K). Therefore, as with experiments using cell pellets, these results suggest that cells may sense inducing signals relative to their neighbours, rather than the absolute local amount of inducing signal.

**Fig. 2.**
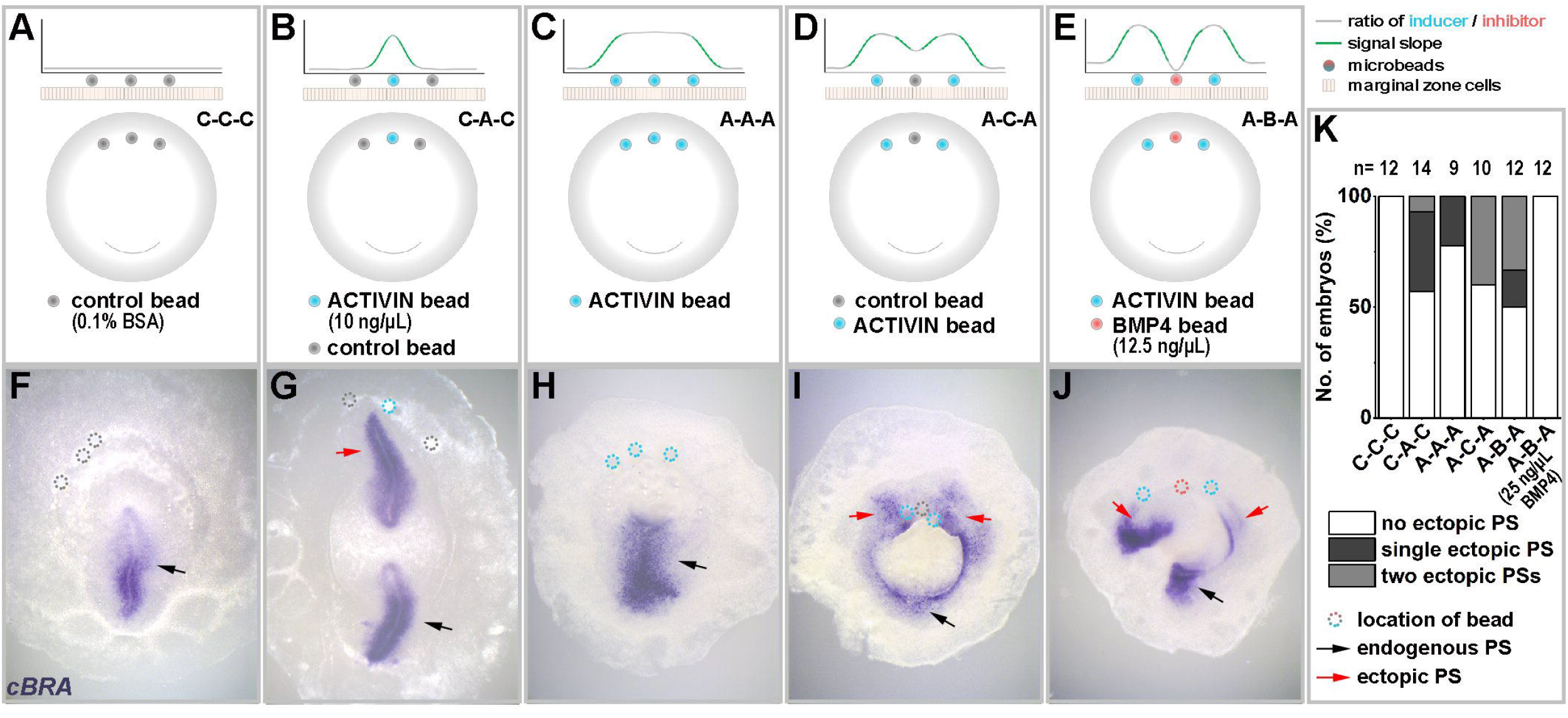
Interruption of a domain exposed to an inducing signal increases the incidence of primitive streak induction – experiments with protein-soaked beads. **(A-E)** Experimental design. Induction of a streak is assessed after five combinations of bead grafts: 3 control beads **(A)**, An ACTIVIN-soaked bead flanked by control beads (B), exposing a wide area to the inducing signal by grafting 3 ACTIVIN-soaked beads **(C)**, interrupting the inducing signal by adding a ‘space’ (control bead) to separate two adjacent inducing (ACTIVIN) beads **(D)**, and adding an inhibitor (BMP4-soaked bead) to separate two adjacent ACTIVIN beads **(E)**. **(F-J)** Representative embryos for each experiment. Two primitive streaks only form when the inducing signal is interrupted, even when adding an inhibitory signal. **(K)** Summary graph showing the incidence of each type of result for the above experiments. Note that a higher concentration of BMP4 (25 ng/μl), does not allow an ectopic streak to form. Dotted circles, location of beads. Other abbreviations and symbols as in Fig. 1.

### Two alternative models

To distinguish between the two alternative mechanisms of how cells might sense their positions (absolute local morphogen concentration or comparison of local signal strength in relation to their neighbourhood), two mathematical models were designed, one for each of these mechanisms, to make experimentally-testable predictions (for details see Materials and Methods). We model the marginal zone as a one-dimensional ring of cells (Fig. 3 A). Positional information is provided by the balance between an inducer (SMAD2/3 activation in response to a VG1/ACTIVIN/NODAL-type signal) and an inhibitor (SMAD 1/5/8 in response to a BMP signal) within each cell (Fig. 3 B). Model A proposes that each cell independently assesses the concentration of morphogens (inducer vs. inhibitor) it receives: when a threshold is exceeded, the cell is triggered to start primitive streak formation. Model B proposes that cells communicate with their neighbours to assess how the streak-inducing signal changes in space: each cell in the ring compares itself with the average signal strength in its neighbourhood to determine whether or not to initiate streak formation (Fig. 3 B).

**Fig. 3.**
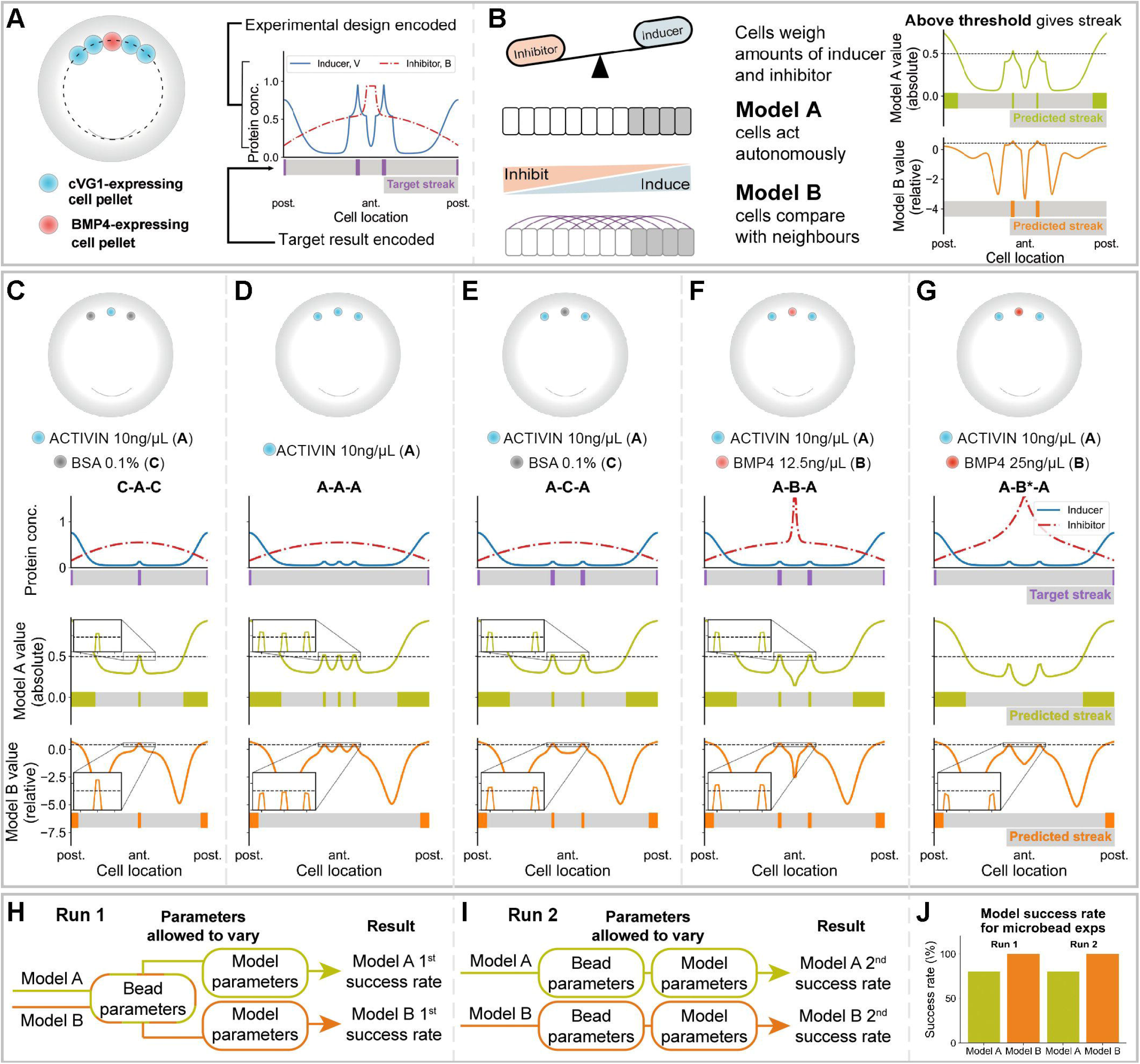
Mathematical model and verification *in silico*. **(A-B)** Model workflow. **(A)** The dotted line represents the marginal zone. Concentrations of primitive-streak-inducing and -inhibiting proteins are inferred from experimental design. Target site of streak initiation is encoded for comparison with model predictions. **(B)** In each cell, both models weigh concentrations of streak-inducing and -inhibiting proteins. Model A assumes that cells act autonomously to define the site of streak formation. Model B assumes that cells compare concentrations within a given neighbourhood to initiate streak formation. Model values are plotted for the entire embryo, where values above a threshold define the site of streak initiation. **(C-G)** *In silico* simulations of bead experiments in Figure 2. Top, experimental designs. First row of plots: inducer levels shown as a red line, inhibitor in blue; the lower bar marks the expected position of streak initiation. Second row of plots: Model A values and corresponding predicted streak locations. Third row: Model B values and streak locations. **(C, E-G)** A model is defined as “successful” for one experimental design if the predicted number and location of streaks matches the target result. **(D)** Model A fails to replicate the experimental result. No parameter values are found where Model A is successful for both designs (C) and (D). **(E-G)** Unlike Model A, Model B predicts that exchanging the control bead for a bead of low dose inhibitor will counter-intuitively increase the chances of ectopic streak formation (insets). **(H-J)** To ensure that finding a single set of parameters does not limit the ability of either model to replicate the target results, we used two approaches for parameter estimation: **(H)** a single set of bead parameters is defined for both models, or **(I)** bead and model parameters vary freely for both models, allowing the maximum chance of success. **(J)** approach H does not reduce the success rate of either model. Model B outperforms Model A in all cases.

As an initial test of the model comparison method, we asked whether there exist parameter values allowing both models to replicate the experimental results shown in Fig. 2. We automated the search for parameter values using Bayesian computation, which scores values with a ‘likelihood function’ (Fig. S3). This function quantifies how well the predicted number and position of ectopic streaks match experimental results on a cell-by-cell basis. All parameter values found were tested for their ability to predict the initiation of ectopic primitive streaks in the appropriate locations in terms of ‘success’ or ‘failure’ of the predictions of each model for each embryonic manipulation. While many parameter values yielded the same model success rates in the 5 experiments illustrated in Fig. 3 C-G (where ‘target streak’ is the experimental result), the likelihood function (Fig. S4) allowed further discrimination. Fig. 3 C-G illustrates the output of each simulation when run with a set of parameter values that provides both the greatest success rate for each model and the highest likelihood score. Even when the parameters were chosen to favour Model A, no set of parameter values was found that allowed Model A to replicate both experimental results in Fig. 3 C, D (the consequences of placing one or three beads of inducer in the anterior marginal zone). In contrast, Model B successfully predicts that broadening the domain of ectopic inducer reduces the chance of initiating ectopic streak formation (Fig. 3 D), even for a set of parameters favouring Model A.

The two models also differ in their ability to portray the effects of placing a bead of inhibitor between two beads of inducer (Fig. 3 E-G). Model A predicts that the presence of the inhibitor will reduce the likelihood of ectopic streaks (Fig. 3 E, F). However, Model B correctly predicts that only low dose of inhibitor increases the chance of forming an ectopic streak (Fig. 3 F, G). The same results were obtained irrespective of whether the sources of inducer and inhibitor were of small diameter (Fig. 3 C-G, to simulate microbeads as in Fig. 2) or wider (Fig. S5, simulating a cell pellet as in Fig. 1).

We sought a single set of bead parameters that would allow both models to mimic the experimental findings (Fig. 3 H). However, choosing a single set of bead parameters could act as a constraint, giving an advantage to one of the models. Therefore, we also performed the parameter inference to allow bead parameters to vary for each model independently (Fig 3 I). Strikingly, Model B always outperforms Model A, regardless of whether a single set of parameters is chosen to fit both models, or whether parameter values are optimised for each model separately (Fig. 3 J).

### Challenging the models and testing predictions

#### a. Decreasing the amount of inhibitor

In both models, cells measure their position by assessing the relative strength of the intracellular downstream effectors of the inducers (VG1/NODAL/ACTIVIN) and inhibitors (BMP). Therefore, decreasing the streak-inhibiting signal alone should induce ectopic primitive streak formation. In this case, both models predict this outcome (Fig. 4 A and B).

**Fig. 4.**
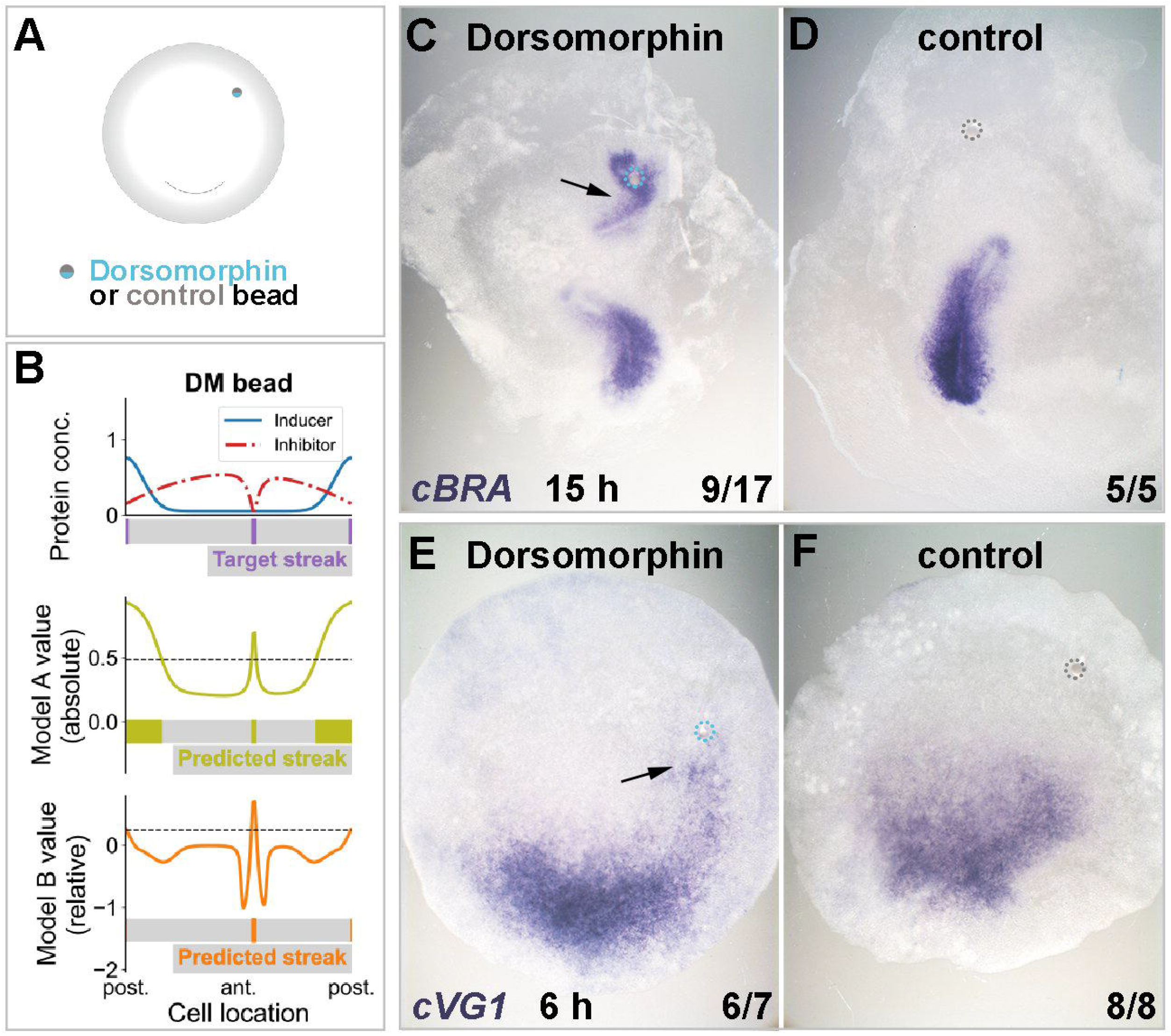
Decreasing the amount of inhibitor induces ectopic primitive streak formation. Local repression of inhibitor (BMP) using Dorsomorphin induces a streak both *in silico* and *in vivo*. **(A)** Experimental setup. **(B)** Results of *in silico* simulations (colours and other conventions as in Fig. 3). Both models predict ectopic primitive streak formation when the concentration of inhibitor is decreased locally. **(C-F)** Results of *in vivo* experiments. A graft of a 1mM Dorsomorphin-soaked bead in the anterior marginal zone induces formation of an ectopic streak expressing *cBRA* after overnight culture (**C**, arrow), which is preceded (at 6 h) by ectopic expression of *cVG1* (**E**, arrow). Control (0.2% DMSO) beads have no effect (**D, F**). Dotted circles, location of microbeads. The proportion of embryos showing the phenotype illustrated are indicated in the lower right of each panel.

To test these predictions experimentally, we used dorsomorphin, an inhibitor of BMP signalling (Yu et al., 2008). A dorsomorphin-soaked bead was grafted in the anterior marginal zone (Fig. 4 A). After overnight culture, an ectopic primitive streak (with *cBRA* expression) was seen to arise close to the bead (Fig. 4 C and D). This result is consistent with a previous study showing that a graft of a cell pellet expressing the BMP antagonist CHORDIN in the area pellucida induces an ectopic streak (Streit et al., 1998). When embryos that had been grafted with a dorsomorphin-bead were examined 6 hours after the graft, ectopic expression of *cVG1* mRNA in the area pellucida (*cVG1* expression is an early target of VG1/NODAL signalling; (Skromne and Stern, 2002, Torlopp et al., 2014)) was found in the vicinity of the bead (Fig. 4 E and F).

#### b. Increasing the amount of inhibitor

A more counterintuitive prediction arises when the strength of inhibition by BMP is increased in a region that normally expresses high levels of BMP (Fig. 5 A). The two models predict different outcomes: Model A predicts that increasing BMP signalling in the anterior marginal zone will reduce the chance of ectopic streak formation (Fig. 5 B). Counterintuitively however, Model B predicts that introducing a bead of inhibitor will increase the streak-inducing values in an area adjacent to the bead (bottom, Fig. 5 B). However Model B also suggests that this effect will be small, perhaps insufficient to result in formation of a mature ectopic primitive streak.

**Fig. 5.**
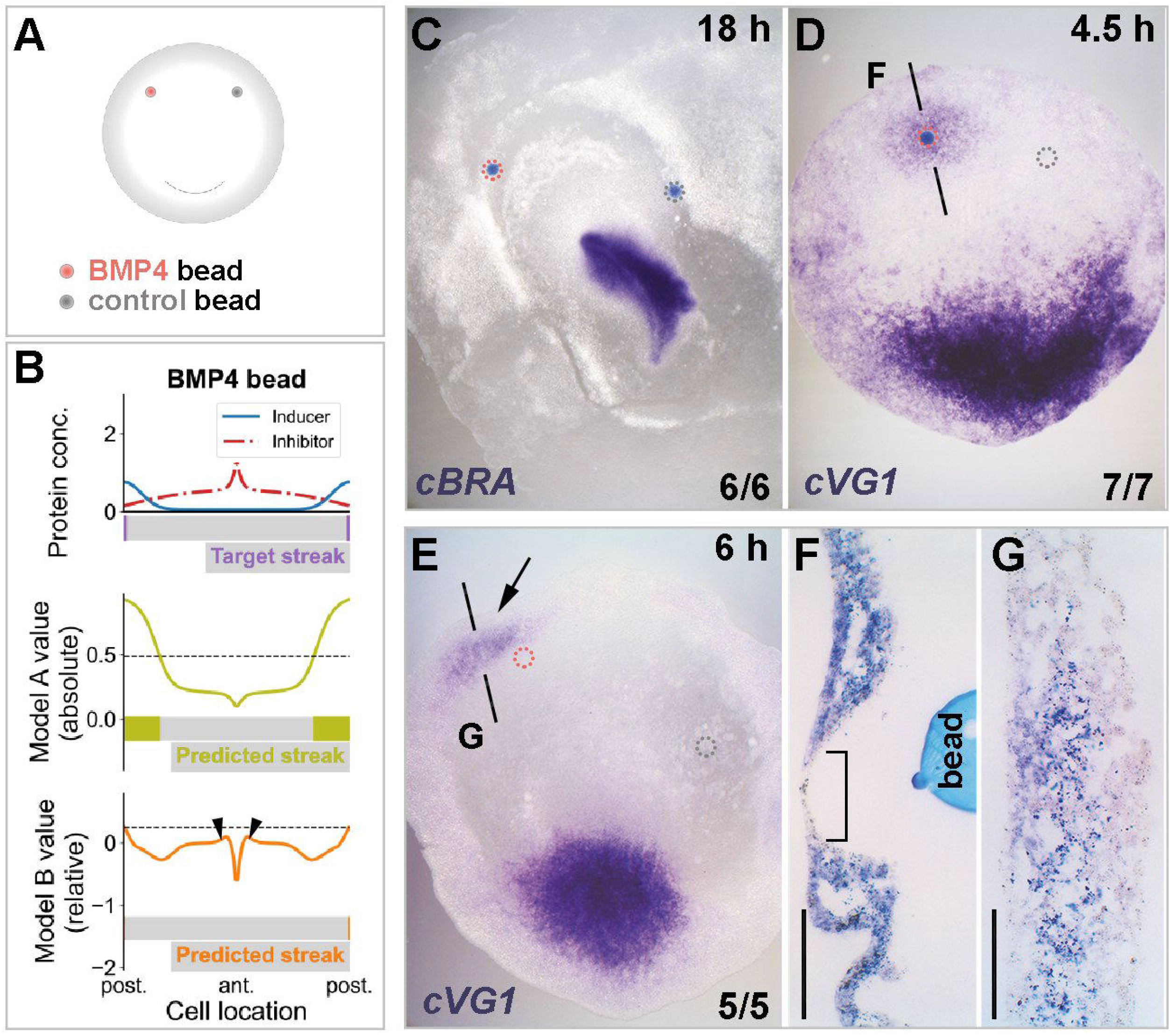
Increasing the amount of inhibitor augments the streak-inducer. Local overexpression of inhibitor (BMP4) increases streak-inducing values *in silico*, and *cVG1* expression *in vivo* in neighbouring cells. **(A)** Experimental setup. **(B)** Results of *in silico* simulations. Only Model B predicts an increase in streak-inducing value in cells neighbouring the bead of inhibitor (arrowheads), but at levels insufficient to initiate an ectopic streak. **(C-G)** Results of *in vivo* experiments. No ectopic primitive streak (marked by *cBRA*) is induced overnight after a graft of BMP4 (50 ng/μl) soaked bead **(C)**. However, a short time (4.5 h) after grafting, ectopic *cVG1* expression is induced in the marginal zone **(D)** in neighbouring cells **(F)** but not in the cells lying directly above the bead (**F**, square bracket). By 6 h after grafting, induced *cVG1* expression is no longer visible in the marginal zone, remaining only in the extraembryonic endoderm (germ wall) (**E**, arrow and **G**). The dashed lines in (**D and E**) indicate the level of the sections in (**F and G**). Dotted circles, location of microbeads. The proportion of embryos showing the illustrated phenotypes is indicated on the lower right of each panel. Scale bar for (**F and G**), 100 μm.

In embryological experiments in which a BMP4 bead was grafted into the anterior marginal zone, no *cBRA* expression or streak formation was observed after overnight incubation (Fig. 5 C). After short incubation (4.5 h), however, *cVG1* expression was observed in cells surrounding the grafted BMP4 bead in the anterior marginal zone and slightly in the adjacent area pellucida (Fig. 5 D). *cVG1*-expression was absent from cells directly overlying the bead (Fig. 5 F) (see also (Arias et al., 2017)). In addition, the ectopic expression was very weak, only detectable after prolonged chromogenic development of the in situ hybridisation (Fig. 5 D and F). This ectopic expression of *cVG1* in the anterior marginal zone was transient: it was seen at 4.5 h and disappeared by 6 h, remaining mostly in the lower layer of the area opaca (germ wall; Fig. 5 E and G). In conclusion, this experimental result conforms with the predictions of Model B but not those of Model A.

#### c. Effect of adjacent sub-threshold amounts of inducer and inhibitor

We have seen that an increase in streak-inhibiting signal can result in paradoxical induction of *cVG1*, which is only predicted by Model B. However, no ectopic *cBRA* expression is observed. If it is indeed the case that cells assess their position in comparison with their neighbours (Model B), rather than measuring the absolute local levels of inducer and inhibitor, then introducing a sub-threshold amount of inducer flanked by low amounts of inhibitor would both deepen and steepen the gradient and might therefore be expected, perhaps paradoxically, to generate a new streak. Model A, in contrast, might predict that neither concentration is high enough locally to affect cell fates resulting in a failure of ectopic streak formation. To simulate this, we explored parameter values for both models that could generate this result (Fig. 6). We find that only Model B can predict the initiation of an ectopic streak (Fig. 6 D-F). No parameters were found that allowed Model A to produce the same result (Fig. 6 D-F).

**Fig. 6.**
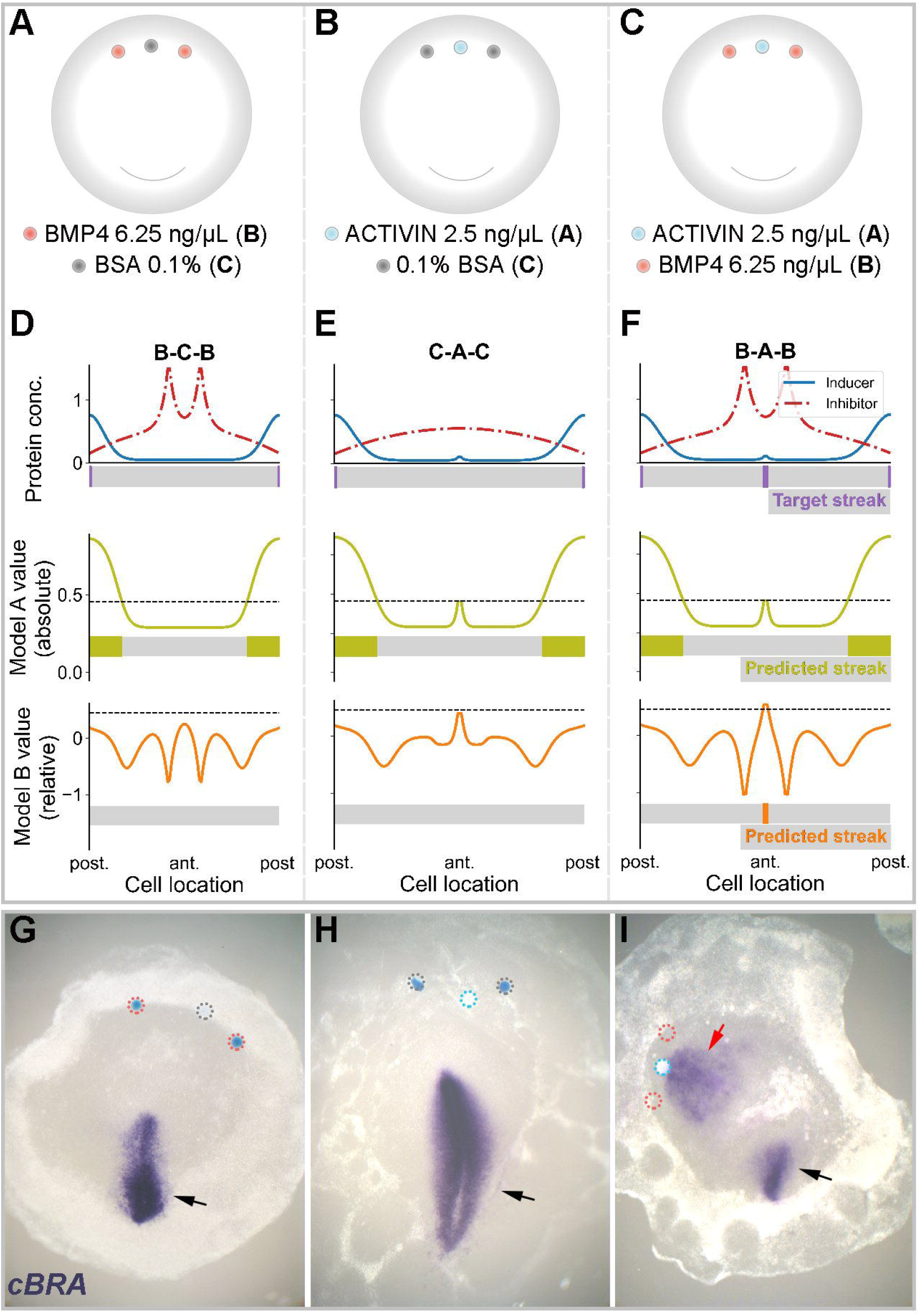
Challenging the models: effect of placing an inhibitor next to sub-threshold amounts of inducer. **(A-C)** Experimental design. Three conditions were tested: two BMP4 beads (6.25 ng/μl) (B) separated by a control bead (C) **(A)**, a bead loaded with sub-threshold (2.5 ng/μl) amounts of ACTIVIN (A) flanked by two control beads (C) **(B)** and a sub-threshold bead of activin flanked by two beads of inhibitor (BMP4) **(C). (D-F)** Results of *in silico* simulations. Only Model B predicts that introducing a sub-threshold amount of inducer flanked by beads of inhibitor will paradoxically generate a site of ectopic PS formation. **(G-I)** Results of *in vitro* experiments showing representative embryos for each experiment. Number of embryos showing the phenotypes are indicated in each panel. *In vivo*, grafting a sub-threshold ACTIVIN bead flanked by two BMP4 beads in the marginal zone can induce ectopic *cBRA* expression (**I**). No such induction is seen in the other combinations (B-C-B or C-A-C) **(A, B, G, H)**. Black and red arrows: endogenous and ectopic *cBRA* expression, respectively. Dotted circles: location of microbeads. The numbers on the lower right of panels G-I indicate the frequency of the illustrated result for each experimental combination.

Next, we tested this prediction experimentally. We began by establishing the minimum threshold of ACTIVIN concentration for PS induction; 2.5 ng/μl of ACTIVIN does not induce cBRA (Fig. S2 D). When two BMP4-beads (6.25 ng/μl) were separated by a control bead, no ectopic PS formed (0/9) (Fig. 6 A and G). When an ACTIVIN-bead (2.5 ng/μl) was flanked by control beads, 97% of embryos showed no ectopic primitive streak (n=37) (Fig. 6 B and H). We then tested the predictions of the model experimentally: when a sub-threshold ACTIVIN bead was flanked by BMP4 beads, *cBRA* expression was seen in 12.5% of cases (n=56) (Fig. 6 C and I). However, a higher concentration of BMP4 (12.5 ng/μl) in the neighbouring beads reduced the proportion of embryos with an ectopic streak (to 9%; n=22) (data not shown), suggesting that at this concentration the total amount of inhibitor may overcome the small amount of inducer emitted by the sub-threshold ACTIVIN-bead. In conclusion, therefore, only Model B correctly predicts the counterintuitive results of this experiment.

Taken together (Fig. 7) our results strongly favour a model by which cells assess their status (in terms of whether or not they will constitute a primitive-streak-initiating centre) in relation to the relative amounts of inducing and inhibiting signals they experience and also in relation to the status of their neighbours, rather than by direct readout of the local concentration of a morphogen that diffuses freely across the entire embryo.

**Fig. 7.**
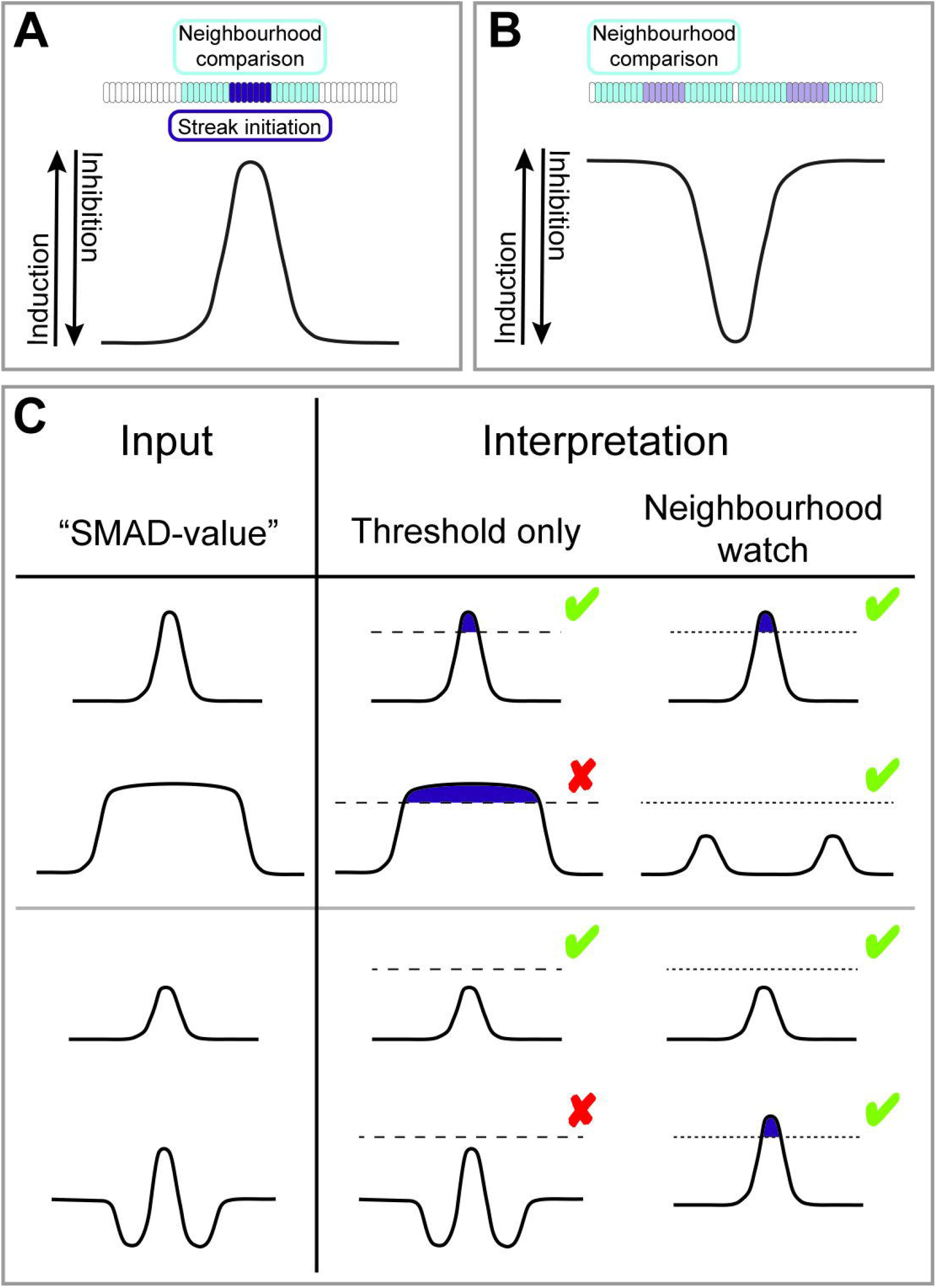
A “neighbourhood watch” model accounts for positioning the site where primitive streak formation is initiated in the marginal zone of the early chick embryo. **(A-B)** The “SMAD-value” represents a combination of inducing and inhibiting signals. Cells assess their position by comparing their SMAD-value with those of their neighbours. Blue: territory over which cells are able to sense. Purple: cell(s) initiating primitive streak formation. Light purple: partial/weak induction. **(A)** The domain of induction must be sufficiently narrow for cells to sense a local maximum. When a local maximum is located, primitive streak formation is initiated in the marginal zone. **(B)** Cells adjacent to a domain of inhibition detect their relatively high SMAD-value and react by emitting streak-inducing signals (cVG1). However, the induction is not sufficiently strong to initiate the formation of a full streak (no *cBRA* expression). **(C)** Comparison of predictions by two models: one (“threshold only”) where positional information is interpreted cell-autonomously solely by assessing the morphogen concentrations, and another (“neighbourhood watch”) where cells make local comparisons with their neighbours to assess their position in the gradients. First row: a narrow domain of induction results in initiation of primitive streak formation. Second row: broadening the domain of induction distinguishes between the two models. The “neighbourhood watch” model predicts that streak formation will not be initiated, matching experimental data. Third row: a sub-threshold amount of inducer results in no ectopic *cBRA* expression. Fourth row, the “threshold only” model predicts that adding inhibitor adjacent to a sub-threshold amount of inducer will either have no effect or reduce the chance of ectopic streak formation. In contrast, the “neighbourhood watch” model correctly predicts the counter-intuitive result that addition of inhibitor increases the chances of ectopic streak initiation. Green ticks and red crosses represent whether the model prediction matches the experimental data or not, respectively. Dashed and dotted lines represent thresholds for interpretation of morphogen concentration. Purple: primitive streak formation initiated in cells above threshold.

## Discussion

Here, we propose a “neighbourhood watch” model to explain how cells interpret positional information to determine the site of gastrulation. Our present results, both from computational modelling and experiments, strongly favour the idea that cells do not read the concentrations of inducer and inhibitor (“SMAD-value”) locally and cell autonomously, but rather interpret their own SMAD-value in relation to that of their neighbours. Moreover, the results suggest that the distance over which such comparisons take place is greater than just the immediately neighbouring cell on either side. In our “neighbourhood watch” model, a neighbourhood size of 100-130 cells is predicted to satisfy experimental observations, based upon the parameter values estimated by the Bayesian inference algorithm.

In previous studies multiple mechanisms have been uncovered by which cells interpret morphogen gradients. Can these other mechanisms explain our results? A key check when answering this question is to ask whether an alternative mechanism can explain the lack of ectopic streak and *cBRA* expression when an inducing signal is applied ectopically as a broad domain (Fig. 2). The first possible mechanism is that cells respond directly to morphogen concentration in a graded manner, as studied in explants of Xenopus embryos with a bead graft (Gurdon et al., 1995). Another study using cultured blastula cells not only supports this but also suggests that interaction with neighbouring cells is not required for the interpretation of morphogen concentration (Gurdon et al., 1999). However, this mechanism cannot explain our result of why a broad domain of inducer paradoxically reduces ectopic *cBRA* expression. A second possible mechanism of morphogen interpretation is one in which cells transform the signal concentration into the intracellular activity of a transcription factor, generating dynamic gene expression patterns with regulatory networks as shown for neural tube patterning (Cohen et al., 2013). Although this mechanism explains well the precision of different thresholds for interpreting morphogen concentrations based on duration and level (strength) of signals, it cannot explain our experimental observations, especially because we find that a broad domain exposed to inducing signal, without changing the duration of signals, reduced the incidence of ectopic *cBRA* expression. These considerations make it more likely that interactions between neighbouring cells are needed to position the primitive streak. A recent paper proposes that a neighbourhood comparison of signal strength (called “spatial fold change (SFC)” model) is required to position the determination front to regulate somite size in the zebrafish trunk and tail bud (Simsek and Ozbudak, 2018), another example of a large developing field undergoing patterning. This suggests that a mechanism involving neighbourhood comparison for the interpretation of positional information may be used by different systems, especially if they are of large size.

In the “neighbourhood watch” model in this study as well as in the SFC model (Simsek and Ozbudak, 2018), cells adopt a relative or normalised value to be evaluated, rather than the absolute morphogen concentration to assess their position. A relative value can provide a stable response of cells to signals, promoting robustness and precision in signal interpretation. Interestingly, a recent *in vitro* study suggests that cells sense relative signal intensity in the TGFβ/SMAD pathway as a fold-change value relative to background to compensate for cellular noise (Frick et al., 2017).

How do cells communicate with their neighbours? In other words, by what mechanism could cells assess their environment? In the wing imaginal disc of Drosophila embryos, the TGFβ-related protein Decapentaplegic (Dpp) acts as a morphogen conveying positional information that results in positioning the wing veins and other features of the wing. Signal-receiving cells have been shown to extend thin and long filopodia, called cytonemes, which extend several cell diameters to the proximity of Dpp-producing cells (Miller et al., 1995, Ramirez-Weber and Kornberg, 1999, Roy et al., 2011). It is worth noting that the existence of filopodia extending very large distances (connecting the invaginating archenteron with the future oral ectoderm at the opposite end of the embryo) was observed by Gustafson and Wolpert in studies of gastrulation in the sea urchin in 1961 (Gustafson and Wolpert, 1961) – this was one of the studies that initiated thinking on pattern formation. Similar structures have been observed in chick embryos during somite development (Sagar et al., 2015) but have not yet been sought at earlier stages. Another important question is: by what mechanism do cells sense relative signals compared to their neighbours? In our simulations, we mimic how each cell encodes the relative strength of inductive (SMAD2/3 activation by Vg1/Nodal/Activin signals) and inhibiting (SMAD1/5/8 activation by BMP) cues they receive as the ratio between them. This is based on the proposal (Candia et al., 1997) that these two classes of SMADs (SMAD2/3 versus SMAD1/5/8) compete for binding to the “co-SMAD”, SMAD4. This could take place in both models - one possible mechanism to provide information about the status of neighbouring cells could involve hypothetical intermediate messengers conveying information about this state. Neighbourhood information could also be transmitted via a positive-feedback mechanism, for example a cell sensing higher levels of BMP would be stimulated to produce more BMP protein (Jones et al., 1992, Metz et al., 1998, Re’em-Kalma et al., 1995, Schulte-Merker et al., 1997).

One question is whether the mechanism proposed here (involving only local cell interactions and no long-range diffusion) is a feature unique to very large fields (several mm), where meaningful positional information conveyed by diffusion alone is likely to be impossible (Crick, 1970). There do appear to be several instances where diffusion of informative morphogens is key, such as initial patterning of the Drosophila blastoderm (Driever and Nusslein-Volhard, 1988a, Gregor et al., 2007b) and mesoderm induction by activin in Xenopus animal caps (Gurdon et al., 1994, Gurdon et al., 1995, McDowell et al., 1997). However, in the chick embryo, the anterior-posterior distance between the two extremes of this ring should span about 300 cell lengths (in reality the marginal zone has a thickness of about 120 μm, corresponding to about 10 cells – here we represent it as being one-cell-thick). As argued by Crick, it seems unlikely that this geometry can support the formation or maintenance of long-range gradients of morphogens generated by free diffusion (Crick, 1970). It therefore seems likely that positional information can be imparted by a variety of different mechanisms, perhaps according to the size and characteristics of the field to be patterned. It will be interesting to perform experiments comparable to those in this paper in a system such as anterior-posterior patterning of the chick limb, which is also large at early stages (HH18-20) and involves a localised signalling region (the Zone of Polarizing Activity) (Riddle et al., 1993, Tickle et al., 1975).

Here we propose that positional information (when interpreted by a collection of cells) defines the location of the signalling centre (NODAL-expressing) that initiates primitive streak formation (Bertocchini and Stern, 2002). Initiation of a streak can be seen as the event that defines embryonic polarity. Our experiments and the associated models were designed to ask questions about how cells within the marginal zone assess their positions around the circumference of this signalling region, and thereafter determine the site next to which (in the area pellucida) the primitive streak will start to form. However, it is important to realise that in the embryo, the downstream consequence of these processes is not only a spot of *cBRA* expression, but rather a true “streak”, gradually extending towards the centre of the embryo. It has been shown previously that this elongation involves a process of cell polarisation and intercalation affecting the same site in the area pellucida where cells receive the inducing signals from the marginal zone (and which itself expresses cVG1 and NODAL) (Rozbicki et al., 2015, Voiculescu et al., 2007, Voiculescu et al., 2014). Here, we observe cases where *cBRA* is induced but this is not followed by formation of an elongated primitive streak. For example, this result is seen when three beads are placed in the anterior marginal zone (A-B-A). One possible reason for this is that the embryos were not incubated for long enough to allow the intercalation to take place, but it is also possible that signals other than cVG1 and inhibition of BMP are required. Indeed it appears that non-canonical (planar cell polarity) WNT signalling drives intercalation (Voiculescu et al., 2007) within the area pellucida. Whatever mechanisms operate in the normal embryo to determine the site of primitive streak formation must somehow coordinate these signalling events to generate the full structure.

Taken together, we provide evidence that in a large system with two opposing gradients, cells assess their position in the field by measuring their location based on the relative concentrations of the inducing (cVG1/NODAL) and inhibitory (BMP) signals, and this is refined by taking cues from their local environment to assess the rate of change of these signals locally. However, the gradients are unlikely to involve long-range diffusion of two morphogens. Regulation of their strength is likely to involve other mechanisms resulting in gradients of transcription and therefore rates of production of the factors.

## Materials and Methods

### Embryo culture and wholemount in situ hybridisation

Fertilised White Leghorn hens’ eggs (Henry Stewart, UK) were incubated for 2-4 hours to obtain EGK X-XI embryos, which were then harvested in Pannett-Compton saline (Pannett and Compton, 1924). After setting up for modified New culture (New, 1955, Stern and Ireland, 1981), the cell pellets or beads were grafted as required for each experiment, and the embryos cultured for the desired length of time before fixation in formaldehyde. Whole mount in situ hybridisation was conducted as previously described (Stern, 1998, Streit and Stern, 2001). The probes used were: *cVG1* (Shah et al., 1997), *cBRA* (Kispert et al., 1995) and *BMP4* (Liem et al., 1995). Stained embryos were imaged under an Olympus SZH10 stereomicroscope with a QImaging Retiga 2000R camera. Some embryos were sectioned in sectioning at 10 μm.

### Misexpression of proteins with transfected cell pellets

HEK293T cells were seeded at 5×10^5^ cells/well in a 6-well dish and incubated for two days (or 1×10^6^ cells/well for transfection on the next day) at 37°C in a total of 2ml 10% FBS DMEM (growth medium)/well. On the day of transfection, the growth medium was changed to 1ml/well of 5% FBS DMEM (transfection medium) at least 30 min before transfection. Transfection was carried out using PEI as reported previously (Papanayotou et al., 2013). Briefly, 3 μl PEI (1mg/ml) was added for every 1 μg of DNA transfected, in a total volume of 150 (for 0.5-2μg)-200μl (for 3-6 μg) DMEM in a sterile Eppendorf. 2μg DNA were transfected/well (containing 6μl PEI/well). Expression plasmids were the previously described DMVg1 (myc-tagged chimeric Vg1 containing the pro-domain of Dorsalin; (Shah et al., 1997), pMT23 (murine BMP4; (Dickinson et al., 1990), and pCAβ-IRES-GFP (as a control). The latter was also used to estimate transfection efficiency. Transfection mixtures were vortexed and then left for 10 minutes at room temperature for the PEI/DNA to complex. The transfection mixture was then added dropwise to the confluent monolayers of cells and incubated overnight at 37°C. The next day cells were checked for transfection efficiency of the GFP plasmid; typically, efficiency ranged from 60-90%. Cells were washed three times with 1 X PBS, trypsinised and resuspended in a total of 1.5ml growth medium and put into a sterile Eppendorf. The cell concentration was estimated in a haemocytometer. A bulk cell suspension of the transfected cells was made in the growth medium, so that each drop contained 500 cells in a total of 20μl growth medium. Hanging drops were formed by placing the 20μl aliquots on the lid of a 6cm cell culture dish, the bottom of which was filled with 5ml of sterile PBS or water to create a humidified atmosphere. After placing several such aliquots well-spaced in a circle, the lid was inverted and placed over the bottom of the dish, creating a mini culture chamber, to allow the cells to coalesce into pellets without adhering to the plastic. Culture dishes were incubated for 36-48 h at 37°C for the formation of pellets ranging in size from 500-1000 cells and used for grafts as required.

### Protein or chemical soaked microbeads

Recombinant human BMP4 (R&D systems, 312-BP) was delivered using Affigel Blue beads (BIO-RAD 1537302); recombinant human ACTIVIN A (R&D systems; 338-AC) was delivered using Heparin-Acrylic beads (Sigma-Aldrich, H5236) and Dorsomorphin hydrochloride (Tocris 3093) was loaded onto AG1X2-formate beads. In each case the beads were incubated overnight at 4 °C in the desired concentration of protein or chemical. Beads were washed in Pannett-Compton saline just before use.

### Encoding the biological problem mathematically

The marginal zone is modelled as a one-dimensional ring of cells, comprising 600 cells in total (based on the assumption that the embryo at this stage contains 20,000-50,000 cells (Bertocchini and Stern, 2012) and on electron microscopy data (Lee et al., 2020, Voiculescu et al., 2007) for estimates of cell size and the radius of the marginal zone). Proxies for streak-inducer and -inhibitor concentrations are assigned to each cell *i*, represented as *V_i_* and *B_i_* respectively with *i* = 1,…,600 (Fig. 3A).

Before the addition of beads, streak-inducer and -inhibitor levels are inferred from a combination of RNAseq reads (Lee et al., 2020) and *in situ* hybridisation of *cVG1* and *cBMP4* (Fig. S1A) respectively, at approximately stage EG&K XII. To mimic these patterns, we use a gaussian function to model the inducer levels based on the observed strong expression of *cVG1* posteriorly, whereas inhibitor levels are modelled with a parabolic function to reflect the shallow, anterior-to-posterior gradient of *cBMP4* (Fig. S1B). The placement of a bead is modelled as having an additive (or subtractive) effect on local protein concentration. The added values are constant for the width of the bead, and then decrease exponentially in space. Therefore, placement of a bead invokes 4 parameters (Fig. S5 A): the position of the centre of the bead, the width of the bead, the bead’s concentration (relating to magnitude of the added values, see Fig. S5 B) and the rate of decay of the added compound in space (i.e. the ‘spread’ parameter of the exponential distribution, see Fig. S5 C).

### Defining two models

For each cell to make its decision to initiate streak formation, we define the relationship between the amounts of SMAD2/3 (as a proxy for amount of inducer received) and SMAD1/5/8 (as a proxy for amount of inhibitor received) within the cells. This is based on the fact that inducing TGFβ-related signals (VG1/ACTIVIN/NODAL) act by phosphorylation of SMAD2/3, whereas inhibitory TGFβ-related signals (BMPs) phosphorylate SMAD1/5/8 – cells have been proposed to evaluate the relative strength of signals through competition of binding of these two classes of SMADs to the “co-SMAD”, SMAD4 (Candia et al., 1997). Inducing and inhibitory SMADs compete to form complexes with a fixed, limited amount of SMAD4. The inducer- and inhibitor-linked SMAD complexes then move to the nucleus and regulate expression of different target genes.

With *V_i_* and *B_i_* representing the levels of inducer- and inhibitor-linked SMAD complexes in cell *i* respectively, we can then represent the total amount of SMAD4 in a cell as the sum of the unbound, inducer-associated and inhibitor-associated SMAD4 (1 + *a_V_V*_i_ + *a_B_B_i_*), where *a_V_* and *a_B_* are scalings of the protein concentrations. We then represent the proportion of streak-inducing SMAD complex in a cell as

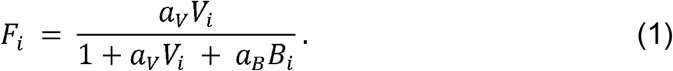

*F_i_* will hereafter be referred to as the “SMAD-value”, with higher values indicating stronger induction.

We define two models for how cells interpret the SMAD-value to make the decision to initiate a primitive streak.

A. Each cell compares its SMAD-values with a fixed threshold, without reference to its neighbours. If the threshold is exceeded, the cell is defined to take part in primitive streak initiation and will express *cBRA*. For each cell *i*, if

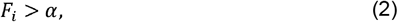

then that cell forms part of the primitive streak initiating domain.
B. Each cell compares its own SMAD-value with those of its neighbours. Each cell can sense these values a certain distance away from itself and calculates an average SMAD-value for all the neighbours it can see. If its own value is sufficiently large compared to the average of its neighbours, the cell becomes part of a primitive streak initiating centre, and expresses *cBRA*. Therefore, a streak is initiated next to cell *i* if

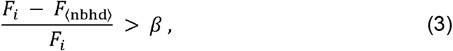

where *F*_〈nbhd〉_ is defined to be the average value of *F_j_* in a given neighbourhood surrounding cell *i*. Specifically,

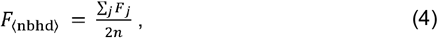

with *j* ∈ [*i* − *n, i* + *n*] \ {*i*}, where (2*n* + 1) is the full width of the neighbourhood.

Both Models A and B have as parameters a threshold value (*α* or *β* and protein concentration scalings (*a_V_* and *a_B_*). Additionally, Model B requires the size of the neighbourhood (*n*) to be defined as a parameter.

### Parameter inference

For the final stage of the modelling process, we ask whether there exists a set of parameters allowing each model to replicate a target result. As both models invoke many parameters, resulting in a large and high-dimensional parameter space, we automate the search with a MCMC Bayesian computation algorithm. Parameter values are scored using a likelihood function which quantifies how well model predictions match a target result. The target result is defined based upon an experimental result (Fig. 3 and Fig. S5) or a new possible theory (Figs. 4–6).

For the parameter search, we fix the expected width of the streak initiating domain, as well as the positions and widths of the beads. We allow the concentration and spread parameters of the beads to vary (denoted *c* and *s*) in addition to all model parameters (*α, β, a_V_, a_B_, n*). Uniform prior distributions are defined for all parameters except the protein concentration scalings, *a_V_* and *a_B_*. For these parameters we define *b_V_* = log_10_ *a_V_* and *b_B_* = log_10_ *a_B_*, which are then uniformly distributed. We define biologically plausible ranges within which parameters are allowed to vary (shown in Fig. S7).

In order to obtain the likelihood function, we first define for each cell, the distance (*f_i_*) of the SMAD-value (*F_i_*) to the threshold for streak formation, which for Model A is

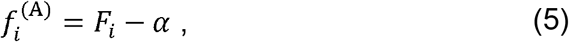

and for Model B

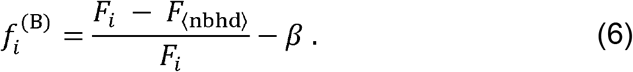

So *f_i_* > 0 implies that a streak will form in cell *i*, and *f_i_* ≤ 0 implies no streak will form. For convenience we can write that *f_i_* = *f_i_*(*θ*) where *θ* = {*α, β, a_V_, a_B_, n, c, s*}, the set of parameters to be varied.

The target result is encoded as a binary decision for each cell: presence or absence of *cBRA* expression indicating the site of primitive streak formation. We therefore define

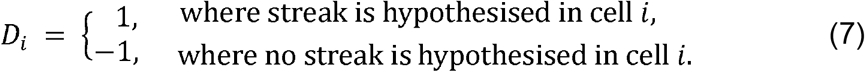

Then the ‘likelihood’ of parameters *θ* can be calculated as

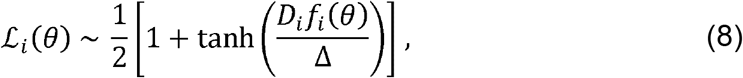

in cell *i*, which approximates a step function as Δ → 0 (Fig. S6). For all parameter searches, we use Δ= 0.05. The likelihood is calculated individually for each cell of each experimental design given to the algorithm. The product of the likelihoods (across cells, designs and parameters) is calculated giving the total likelihood for a given set of parameter values. The parameters used to calculate the total likelihood include all model parameters and the bead parameters relevant for the experiment. Only cells in the anterior half of the embryo are used to calculate the total likelihood, because beads are only grafted anteriorly in the experiments modelled. As a result of this, Model B does not always predict the presence of an endogenous streak next to the posterior margin.

The posterior distributions of the parameters were obtained via the MCMC Bayesian computation in the pyDREAM package (Shockley et al., 2017) which implements a DREAM_(ZS)_ algorithm (Laloy and Vrugt, 2012). The algorithm was run using 5 Markov chains for a minimum of 5000 iterations per chain, and convergence was tested using the Gelman–Rubin statistic (Gelman and Rubin, 1992, Brooks and Gelman, 1998). The posterior distributions are shown in Figure S7. An approximate neighbourhood size can be inferred from the posterior distribution of the parameter *n* (defined in equation 4), which peaks between 50-65 cells for all experiments.

The Bayesian computation algorithm maximises the likelihood (equation 8), quickly and efficiently finding sets of parameter values minimizing the distance between the target result (*D_i_*) and the model result (*f_i_*). Specifically, the likelihood function is defined so as to strongly favour sets of parameters where *D_i_* and *f_i_* have the same sign (i.e. both above zero or both below zero). Occasionally this means that parameter values obtained by the algorithm give model values close to, but not exceeding, the threshold and therefore do not predict ectopic streaks as required by the target result. Therefore, all parameter values found using the Bayesian computation algorithm were checked to the ensure that ectopic streaks were predicted in locations dictated by the target result. This was done by verifying that at least one cell exceeded the threshold to produce an ectopic streak in the expected location (i.e. the location of a bead). Thus, if parameter values for a given model allowed the prediction of correct ectopic streak placement, these values were deemed to give ‘success’ for a specific experimental design. The parameter values used in the plots in Figures 3–6 and S3 were chosen to maximise both the success rate and the likelihood. We have verified that there is a positive correlation between the success rate and the likelihood score (Fig. S4). All parameter values are given in Data S1.

The parameter search is performed for each group of experimental designs comprising Figures 1, 2, 4/5 and 6. Ideally, the parameter search must output a single set of bead parameters, allowing both models to approximate the target results as closely as possible (Fig. 3 H). However, this acts as a restriction that might limit the ability of either model to replicate the target result. Therefore, the parameter search was also performed with all parameters varying for both models independently removing this restriction (Fig. 3 I). We verified that seeking a single set of bead parameters did not reduce the ability of either model to replicate the target result (Fig. 3 J).

## Supporting information

Supplemental information including figures

## Author contributions

HCL conducted all embryo experiments; CH designed the models and implemented them; NMMO constructed the vectors and performed cell culture; RPC and KP provided advice and ideas on mathematical methods; LW provided inspiration and stimulated questions during the early stages of the study; CDS supervised the study. HCL, CH and CDS wrote the paper.

## Data and materials availability

The software used for the mathematical and computational modelling is available at https://github.com/catohaste/neighbourhood-streak.

## References

Arias, C. F., Herrero, M. A., Stern, C. D. & Bertocchini, F. 2017. A molecular mechanism of symmetry breaking in the early chick embryo. Sci Rep, 7, 15776.

Bertocchini, F., Skromne, I., Wolpert, L. & Stern, C. D. 2004. Determination of embryonic polarity in a regulative system: evidence for endogenous inhibitors acting sequentially during primitive streak formation in the chick embryo. Development, 131, 3381–90.

Bertocchini, F. & Stern, C. D. 2002. The hypoblast of the chick embryo positions the primitive streak by antagonizing nodal signaling. Dev Cell, 3, 735–44.

Bertocchini, F. & Stern, C. D. 2012. Gata2 provides an early anterior bias and uncovers a global positioning system for polarity in the amniote embryo. Development, 139, 4232–8.

Bode, H. 2011. Axis formation in hydra. Annu Rev Genet, 45, 105–17.

Brooks, S. P. & Gelman, A. 1998. General Methods for Monitoring Convergence of Iterative Simulations. Journal of Computational and Graphical Statistics, 7, 434–455.

Candia, A. F., Watabe, T., Hawley, S. H., Onichtchouk, D., Zhang, Y., Derynck, R., Niehrs, C. & Cho, K. W. 1997. Cellular interpretation of multiple TGF-beta signals: intracellular antagonism between activin/BVg1 and BMP-2/4 signaling mediated by Smads. Development, 124, 4467–80.

Cohen, M., Briscoe, J. & Blassberg, R. 2013. Morphogen interpretation: the transcriptional logic of neural tube patterning. Curr Opin Genet Dev, 23, 423–8.

Crick, F. 1970. Diffusion in embryogenesis. Nature, 225, 420–2.

Dessaud, E., Ribes, V., Balaskas, N., Yang, L. L., Pierani, A., Kicheva, A., Novitch, B. G., Briscoe, J. & Sasai, N. 2010. Dynamic assignment and maintenance of positional identity in the ventral neural tube by the morphogen sonic hedgehog. PLoS Biol, 8, e1000382.

Dickinson, M. E., Kobrin, M. S., Silan, C. M., Kingsley, D. M., Justice, M. J., Miller, D. A., Ceci, J. D., Lock, L. F., Lee, A., Buchberg, A. M. & ET AL. 1990. Chromosomal localization of seven members of the murine TGF-beta superfamily suggests close linkage to several morphogenetic mutant loci. Genomics, 6, 505–20.

Driever, W. & Nusslein-Volhard, C. 1988a. The bicoid protein determines position in the Drosophila embryo in a concentration-dependent manner. Cell, 54, 95–104.

Driever, W. & Nusslein-Volhard, C. 1988b. A gradient of bicoid protein in Drosophila embryos. Cell, 54, 83–93.

Frick, C. L., Yarka, C., Nunns, H. & Goentoro, L. 2017. Sensing relative signal in the Tgf-beta/Smad pathway. Proc Natl Acad Sci U S A, 114, E2975–E2982.

Gelman, A. & Rubin, D. B. 1992. Inference from Iterative Simulation Using Multiple Sequences. Statistical Science, 7, 457–472.

Green, J. B. & Smith, J. C. 1990. Graded changes in dose of a Xenopus activin A homologue elicit stepwise transitions in embryonic cell fate. Nature, 347, 391–4.

Gregor, T., Tank, D. W., Wieschaus, E. F. & Bialek, W. 2007a. Probing the limits to positional information. Cell, 130, 153–64.

Gregor, T., Wieschaus, E. F., Mcgregor, A. P., Bialek, W. & Tank, D. W. 2007b. Stability and nuclear dynamics of the bicoid morphogen gradient. Cell, 130, 141–52.

Gurdon, J. B., Harger, P., Mitchell, A. & Lemaire, P. 1994. Activin signalling and response to a morphogen gradient. Nature, 371, 487–92.

Gurdon, J. B., Mitchell, A. & Mahony, D. 1995. Direct and continuous assessment by cells of their position in a morphogen gradient. Nature, 376, 520–1.

Gurdon, J. B., Standley, H., Dyson, S., Butler, K., Langon, T., Ryan, K., Stennard, F., Shimizu, K. & Zorn, A. 1999. Single cells can sense their position in a morphogen gradient. Development, 126, 5309–17.

Gustafson, T. & Wolpert, L. 1961. Studies on the cellular basis of morphogenesis in the sea urchin embryo. Gastrulation in vegetalized larvae. Exp Cell Res, 22, 437–49.

Jones, C. M., Lyons, K. M., Lapan, P. M., Wright, C. V. & Hogan, B. L. 1992. DVR-4 (bone morphogenetic protein-4) as a posterior-ventralizing factor in Xenopus mesoderm induction. Development, 115, 639–47.

Kispert, A., Ortner, H., Cooke, J. & Herrmann, B. G. 1995. The chick Brachyury gene: developmental expression pattern and response to axial induction by localized activin. Dev Biol, 168, 406–15.

Kumar, A., Gates, P. B. & Brockes, J. P. 2007. Positional identity of adult stem cells in salamander limb regeneration. C R Biol, 330, 485–90.

Laloy, E. & Vrugt, J. A. 2012. High-dimensional posterior exploration of hydrologic models using multiple-try DREAM(ZS) and high-performance computing. Water Resources Research, 48.

Lecuit, T., Brook, W. J., Ng, M., Calleja, M., Sun, H. & Cohen, S. M. 1996. Two distinct mechanisms for long-range patterning by Decapentaplegic in the Drosophila wing. Nature, 381, 387–393.

Lee, H. C., Lu, H. C., Turmaine, M., Oliveira, N. M. M., Yang, Y., De Almeida, I. & Stern, C. D. 2020. Molecular anatomy of the pre-primitive-streak chick embryo. Open Biol, 10, 190299.

Liem, K. F., JR., Tremml, G., Roelink, H. & Jessell, T. M. 1995. Dorsal differentiation of neural plate cells induced by BMP-mediated signals from epidermal ectoderm. Cell, 82, 969–79.

Mcdowell, N., Zorn, A. M., Crease, D. J. & Gurdon, J. B. 1997. Activin has direct long-range signalling activity and can form a concentration gradient by diffusion. Curr Biol, 7, 671–81.

Metz, A., Knochel, S., Buchler, P., Koster, M. & Knochel, W. 1998. Structural and functional analysis of the BMP-4 promoter in early embryos of Xenopus laevis. Mech Dev, 74, 29–39.

Miller, J., Fraser, S. E. & Mcclay, D. 1995. Dynamics of thin filopodia during sea urchin gastrulation. Development, 121, 2501–11.

Mitrani, E., Ziv, T., Thomsen, G., Shimoni, Y., Melton, D. A. & Bril, A. 1990. Activin can induce the formation of axial structures and is expressed in the hypoblast of the chick. Cell, 63, 495–501.

New, D. A. T. 1955. A new technique for the cultivation of the chick embryo in vitro. J. Embryol. exp. Morph., 3, 326–331.

Pannett, C. A. & Compton, A. 1924. The cultivation of tissues in saline embryonic juice. Lancet, 1, 381–384.

Papanayotou, C., De Almeida, I., Liao, P., Oliveira, N. M., Lu, S. Q., Kougioumtzidou, E., Zhu, L., Shaw, A., Sheng, G., Streit, A., Yu, D., Wah Soong, T. & Stern, C. D. 2013. Calfacilitin is a calcium channel modulator essential for initiation of neural plate development. Nat Commun, 4, 1837.

Postlethwait, J. H. & Schneiderman, H. A. 1971. Pattern formation and determination in the antenna of the homoeotic mutant Antennapedia of Drosophila melanogaster. Dev Biol, 25, 606–40.

Ramirez-Weber, F. A. & Kornberg, T. B. 1999. Cytonemes: cellular processes that project to the principal signaling center in Drosophila imaginal discs. Cell, 97, 599–607.

Re’em-Kalma, Y., Lamb, T. & Frank, D. 1995. Competition between noggin and bone morphogenetic protein 4 activities may regulate dorsalization during Xenopus development. Proc Natl Acad Sci U S A, 92, 12141–5.

Riddle, R. D., Johnson, R. L., Laufer, E. & Tabin, C. 1993. Sonic hedgehog mediates the polarizing activity of the ZPA. Cell, 75, 1401–16.

Roy, S., Hsiung, F. & Kornberg, T. B. 2011. Specificity of Drosophila cytonemes for distinct signaling pathways. Science, 332, 354–8.

Rozbicki, E., Chuai, M., Karjalainen, A. I., Song, F., Sang, H. M., Martin, R., Knolker, H. J., Macdonald, M. P. & Weijer, C. J. 2015. Myosin-II-mediated cell shape changes and cell intercalation contribute to primitive streak formation. Nat Cell Biol, 17, 397–408.

Sagar, Prols, F., Wiegreffe, C. & Scaal, M. 2015. Communication between distant epithelial cells by filopodia-like protrusions during embryonic development. Development, 142, 665–71.

Schaller, H. C. 1973. Isolation and characterization of a low-molecular-weight substance activating head and bud formation in hydra. J Embryol Exp Morphol, 29, 27–38.

Schulte-Merker, S., Lee, K. J., Mcmahon, A. P. & Hammerschmidt, M. 1997. The zebrafish organizer requires chordino. Nature, 387, 862–3.

Shah, S. B., Skromne, I., Hume, C. R., Kessler, D. S., Lee, K. J., Stern, C. D. & Dodd, J. 1997. Misexpression of chick Vg1 in the marginal zone induces primitive streak formation. Development, 124, 5127–38.

Shockley, E. M., Vrugt, J. A. & Lopez, C. F. 2017. PyDREAM: high-dimensional parameter inference for biological models in python. Bioinformatics, 34, 695–697.

Simsek, M. F. & Ozbudak, E. M. 2018. Spatial Fold Change of FGF Signaling Encodes Positional Information for Segmental Determination in Zebrafish. Cell Rep, 24, 66–78 e8.

Skromne, I. & Stern, C. D. 2002. A hierarchy of gene expression accompanying induction of the primitive streak by Vg1 in the chick embryo. Mech Dev, 114, 115–8.

Stern, C. D. 1998. Detection of multiple gene products simultaneously by in situ hybridization and immunohistochemistry in whole mounts of avian embryos. Curr Top Dev Biol, 36, 223–243.

Stern, C. D. & Ireland, G. W. 1981. An integrated experimental study of endoderm formation in avian embryos. Anat Embryol, 163, 245–63.

Stern, C. D., Yu, R. T., Kakizuka, A., Kintner, C. R., Mathews, L. S., Vale, W. W., Evans, R. M. & Umesono, K. 1995. Activin and its receptors during gastrulation and the later phases of mesoderm development in the chick embryo. Dev Biol, 172, 192–205.

Streit, A., Lee, K. J., Woo, I., Roberts, C., Jessell, T. M. & Stern, C. D. 1998. Chordin regulates primitive streak development and the stability of induced neural cells, but is not sufficient for neural induction in the chick embryo. Development, 125, 507–19.

Streit, A. & Stern, C. D. 1999. Mesoderm patterning and somite formation during node regression: differential effects of chordin and noggin. Mech Dev, 85, 85–96.

Streit, A. & Stern, C. D. 2001. Combined whole-mount in situ hybridization and immunohistochemistry in avian embryos. Methods, 23, 339–44.

Tickle, C., Summerbell, D. & Wolpert, L. 1975. Positional signalling and specification of digits in chick limb morphogenesis. Nature, 254, 199–202.

Torlopp, A., Khan, M. A., Oliveira, N. M., Lekk, I., Soto-Jimenez, L. M., Sosinsky, A. & Stern, C. D. 2014. The transcription factor Pitx2 positions the embryonic axis and regulates twinning. Elife, 3, e03743.

Voiculescu, O., Bertocchini, F., Wolpert, L., Keller, R. E. & Stern, C. D. 2007. The amniote primitive streak is defined by epithelial cell intercalation before gastrulation. Nature, 449, 1049–52.

Voiculescu, O., Bodenstein, L., Lau, I. J. & Stern, C. D. 2014. Local cell interactions and self-amplifying individual cell ingression drive amniote gastrulation. Elife, 3, e01817.

Wolpert, L. 1968. The French flag problem: a contribution to the discussion on pattern development and regeneration. In: Waddington, C. H. (ed.) Towards a Theoretical Biology. Aldine Publishing Co.

Wolpert, L. 1969. Positional information and the spatial pattern of cellular differentiation. Journal of Theoretical Biology, 25, 1–47.

Yu, P. B., Hong, C. C., Sachidanandan, C., Babitt, J. L., Deng, D. Y., Hoyng, S. A., Lin, H. Y., Bloch, K. D. & Peterson, R. T. 2008. Dorsomorphin inhibits BMP signals required for embryogenesis and iron metabolism. Nat Chem Biol, 4, 33–41.

