## Supplemental information including figures for "“Neighbourhood watch” model: embryonic epiblast cells assess positional information in relation to their neighbours"

**Fig. S1. Expression of *cVG1* and *BMP4* in the pre-primitive-streak chick embryo.** **(A)** *cVG1* and *BMP4* are expressed as opposing gradients – *cVG1* is expressed as a very steep gradient decreasing from the posterior marginal zone, and *BMP4* forms a shallower gradient decreasing in a posterior direction. **(B)** Gaussian and parabolic functions are used to model the opposing gradients of inducer and inhibitor, relating to cVG1 and BMP4 respectively.

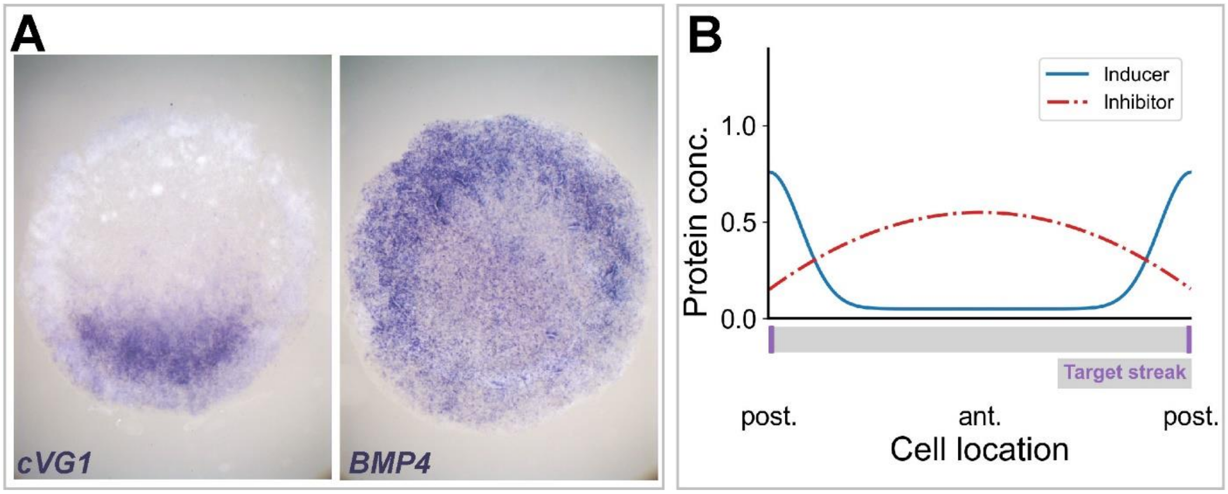

**Fig. S2. Inducing or inhibitory effect of grafting ACTIVIN- or BMP4-soaked microbeads. (A-E)** A graft of an ACTIVIN-soaked bead in the anterior marginal zone induces an ectopic primitive streak (arrow) at concentration of 10 ng/µl **(B)** and 5 ng/µl **(C)**, but not at 2.5 ng/µl **(D)**; **E** shows a control (0.1 % BSA-soaked bead). Dotted circles, position of the bead. The proportion of embryos showing the effect illustrated is indicated in each panel. *cBRA*: primitive streak marker. **(F-J)** A graft of a BMP4-soaked bead in the posterior marginal zone inhibits streak formation. **(J)** summarises the incidence of the various types of result.

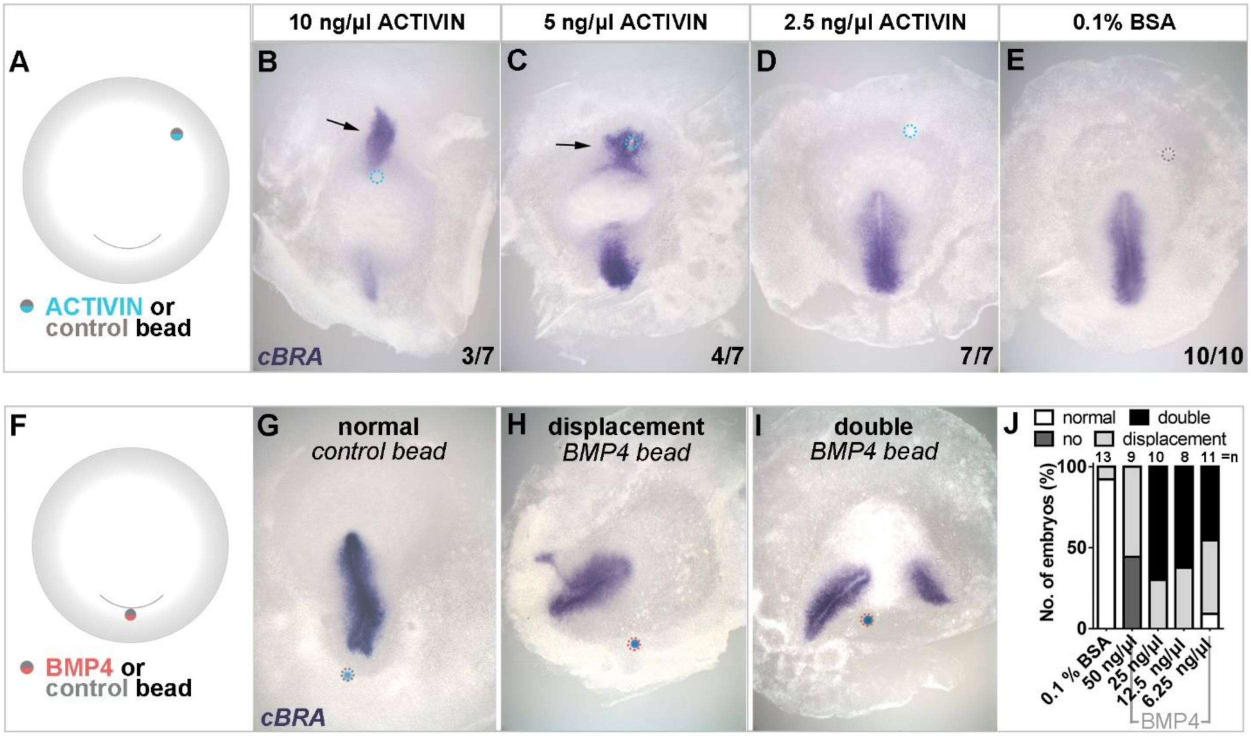

**Fig. S3. Likelihood function used in Bayesian inference of parameters.** The likelihood function is defined so that when a set of parameters allows a model prediction to mimic the target result, the value of the likelihood function is high (and vice-versa). The likelihood function approximates a step function.

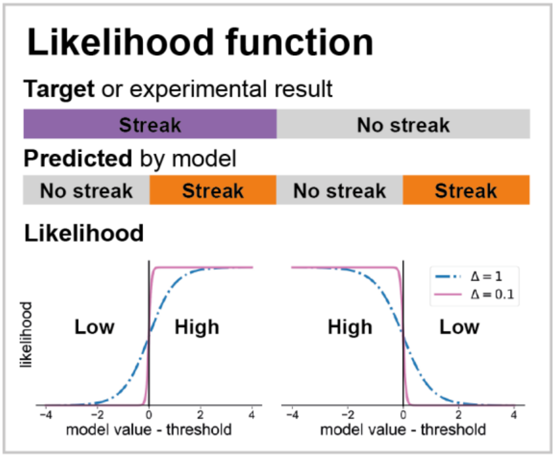

**Fig. S4. Correlation between likelihood and success rate.** The Bayesian parameter inference algorithm samples sets of parameters and assigns them a “likelihood” score, quantifying how well a model can replicate the experimental results. Then, if a set of parameters allows a model to predict the correct number and location of ectopic streaks, the prediction is deemed successful for a single embryonic manipulation. For Figure 3, sets of parameters for each model can be given a “success rate” for all 5 embryonic manipulations shown (Fig. 3 C-G). The likelihood of each parameter set is plotted against its success rate, showing a positive correlation. The top row shows the full range of likelihood scores, while the bottom row shows a narrow subset of this range. For Model A, no set of parameter values was found that gives a 100% success rate. In contrast, multiple sets of parameters were found allowing Model B to predict the experimental results correctly. Here, the parameter estimation was performed with the bead parameters varying separately for each model, giving both models the maximum chance of success, labelled “run 2” in Figure 3J.

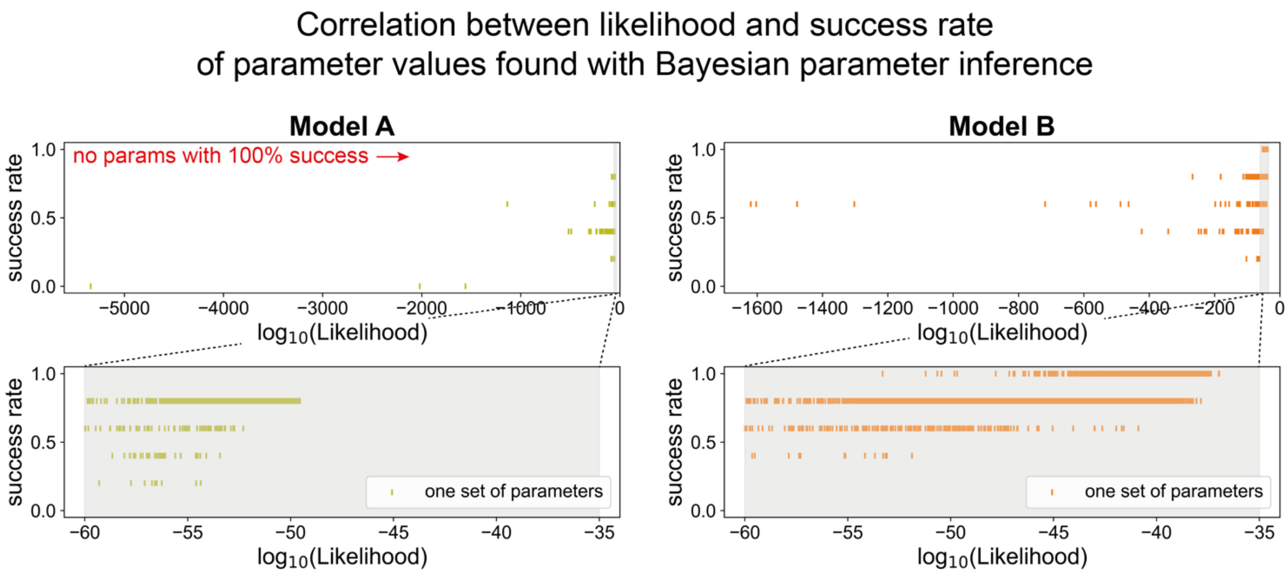

**Fig. S5. *In silico* simulations of cell pellet experiments shown in Fig. 1.** Top row: experimental designs. The first row of plots shows the encoded experimental design and the target result (based on the experimental findings). The second row of plots shows model A values and the corresponding predicted site of primitive streak initiation. The bottom row shows the results obtained with model B. **(A-B)** Model A predicts that broadening the domain of ectopic inducer increases the chance of ectopic *cBRA* expression, whereas model B predicts that the occurrence of ectopic *cBRA* expression will be reduced. The prediction of Model B aligns better with experimental results (Fig. 1 A-B, G, J, M). **(C-D)** Simulated results when a control or BMP4-expressing cell pellet is flanked by cVG1-expressing pellets.

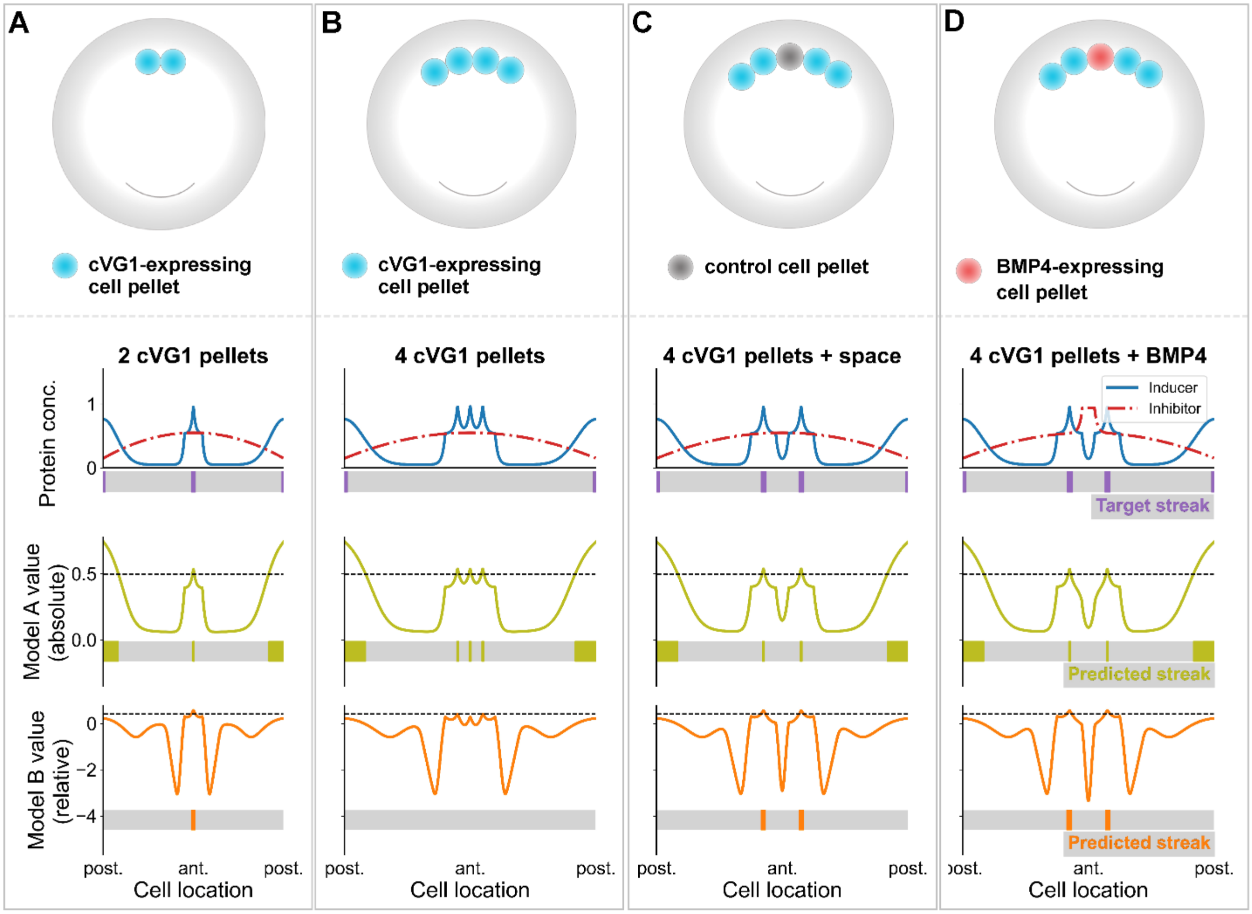

**Fig. S6. Modelling the placement of beads. (A)** The position of the bead relative to the embryo is encoded by the center of the bead and is governed by the experimental design. The placing of a bead causes a constant, additive change in protein concentration throughout the width of the bead. The magnitude of this change is defined as the concentration of the bound protein, *c*. The protein concentration then decays exponentially in space, at a rate governed by the spread parameter, *s*. During parameter estimation, the center and width of the bead are kept constant while the parameters *c* and *s* are permitted to vary. **(B)** Changing the concentration parameter (*c*) principally changes the height of the peak (or trough) and has little effect on the number of cells affected by the bead placement. **(C)** Changing the spread parameter (*s*) has no effect on the magnitude of protein concentration change, but can have a large impact on the size of the territory around the bead that is affected by the ligand.

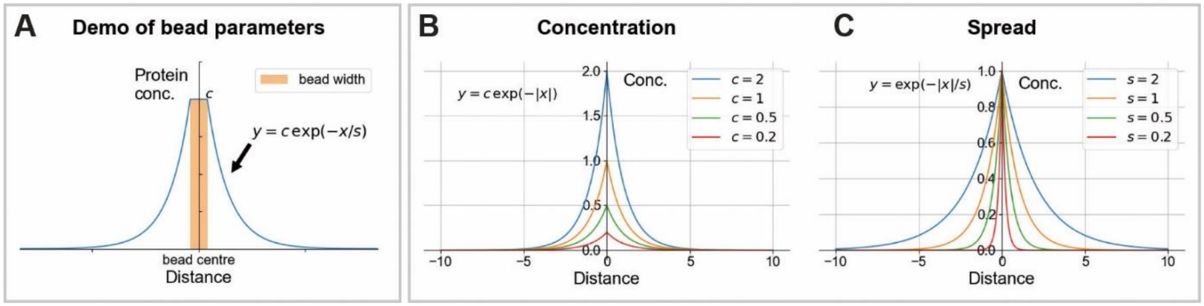

**Fig. S7. Posterior distributions of parameters following Bayesian inference.** Each cell of the table shows posterior distributions resulting from a single parameter search, corresponding to the figures shown. Separate distributions are given for models A and B, as the bead parameters were varied independently for the two models as described in Figure 3 I. In a single cell of the table, the plots along the leading diagonal give the marginal posterior distributions for each parameter. The plots in the top right corner and bottom left corner represent both represent joint distributions for two parameters, allowing the reading to study cross-talk between parameters. In the top right corner, joint distributions are represented with a scatter plot, where each point corresponds to a set of parameters tried during the parameter search and its colour corresponds to the likelihood of this set of parameters. In the bottom left corner, joint distributions are represented by a contour plot where darker colours represent a higher density of parameter values sampled.

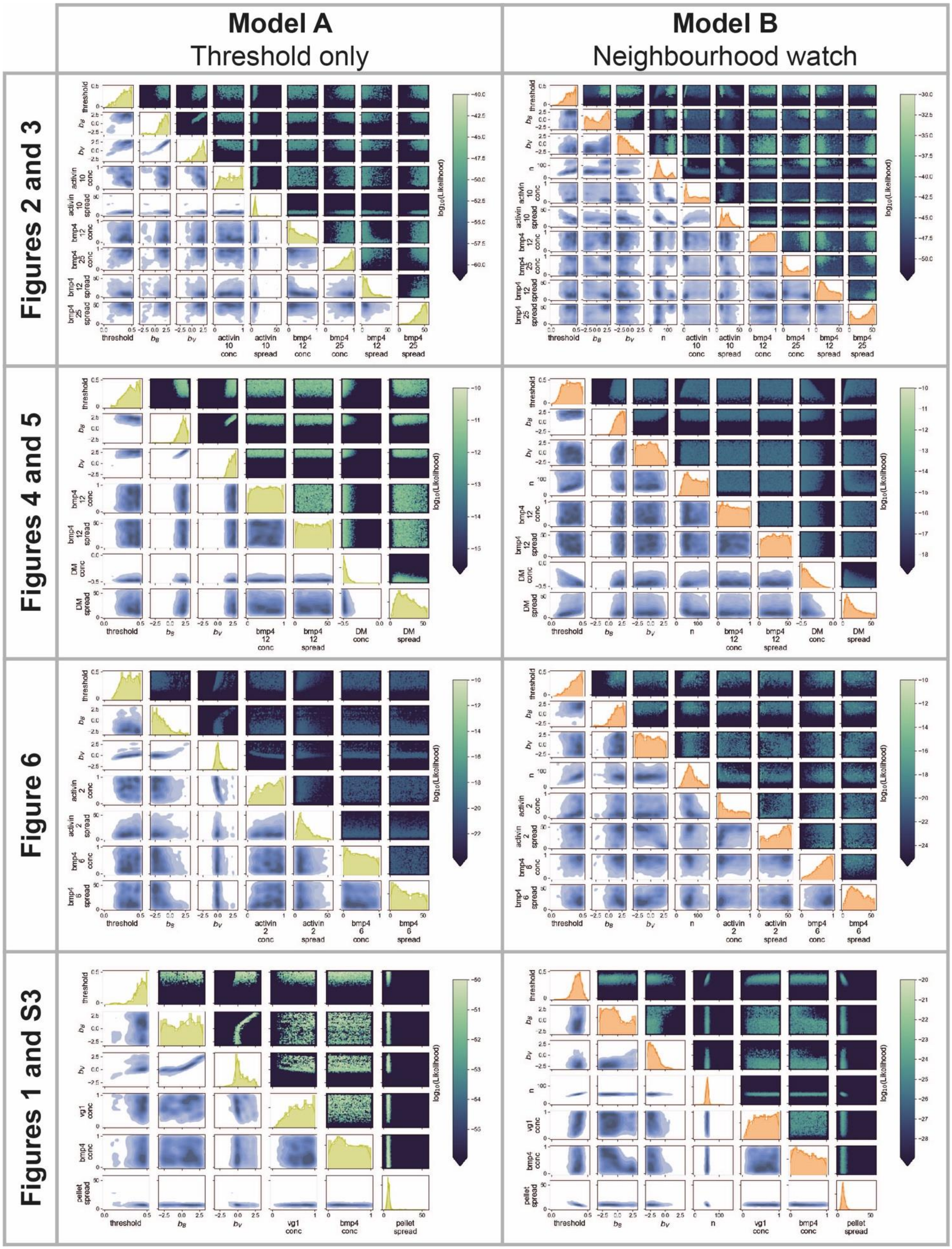

Model A
Threshold only
Model B
Neighbourhood watch
Figures 2 and 3
Figures 4 and 5
Figure 6
Figures 1 and S3
log10(Likelihood)

**Data S1. Results of parameter searches**. The parameter values giving the highest success rate for each parameter search are shown. Parameter searches were run separately for each group of experimental designs (rows 4-15). For each set of experimental designs, models A and B were run separately to give each model the best chance of predicting the experimental results accurately (rows 4-5, 7-8, 10-11, 13-14). The parameter search was also run so as to predict a single set of bead parameters for both models (rows 6, 9, 12, 15). The parameters given are for the pyDREAM algorithm (columns D-F), for each model (columns G-M) and for the type of bead used (columns N-AB). The result of each parameter search is denoted by the total likelihood (column AC), success (TRUE) or failure (FALSE) of each experimental design (columns AD-BE) and the overall success rate of each model (columns BF-BG).
